# CD127 imprints functional heterogeneity to diversify monocyte responses in human inflammatory diseases

**DOI:** 10.1101/2020.11.10.376277

**Authors:** Bin Zhang, Yuan Zhang, Lei Xiong, Yuzhe Li, Yunliang Zhang, Jiuliang Zhao, Hui Jiang, Can Li, Yunqi Liu, Xindong Liu, Haofei Liu, Yi-Fang Ping, Qiangfeng Cliff Zhang, Zheng Zhang, Xiu-Wu Bian, Yan Zhao, Xiaoyu Hu

**Affiliations:** Institute for Immunology and School of Medicine, Tsinghua University, Beijing, China; Beijing Key Laboratory for Immunological Research on Chronic Diseases, Beijing, China; MOE Key Laboratory of Bioinformatics, Beijing Advanced Innovation Center for Structural Biology, Center for Synthetic and Systems Biology, School of Life Sciences, Tsinghua University, Beijing, China; Academy for Advanced Interdisciplinary Studies, Peking University, Beijing, China; Department of Rheumatology, Peking Union Medical College Hospital, Chinese Academy of Medical Science and Peking Union Medical College, National Clinical Research Center for Dermatologic and Immunologic Diseases (NCRC-DID), Beijing, China; Institute of Pathology, Southwest Hospital, Third Military Medical University (Army Medical University), Chongqing, China; Tsinghua-Peking Center for Life Sciences, Beijing, China; Institute for Hepatology, National Clinical Research Center for Infectious Disease, Shenzhen Third People’s Hospital, Shenzhen, China; The Second Affiliated Hospital, School of Medicine, Southern University of Science and Technology, Shenzhen, China; Department of Pathology, Immunology and Laboratory Medicine, University of Florida, Gainesville, Florida, USA; Laboratory of Viral Diseases, National Institute of Allergy and Infectious Diseases, National Institutes of Health, Bethesda, Maryland, USA; Department of Cardiology, National Center for Cardiovascular Diseases Fuwai Hospital, Chinese Academy of Medical Sciences and Peking Union Medical College, Beijing, China

**Author notes:** These authors contributed equally (Bin Zhang, Yuan Zhang).

## Abstract

Studies on human monocytes historically focused on characterization of bulk responses, whereas functional heterogeneity is largely unknown. Here, we identified an inducible population of CD127-expressing human monocytes under inflammatory conditions and named the subset M127. M127 is nearly absent in healthy individuals yet abundantly present in patients with infectious and inflammatory conditions such as COVID-19 and rheumatoid arthritis. Multiple genomic and functional approaches revealed unique gene signatures of M127 and unified anti-inflammatory properties imposed by the CD127-STAT5 axis. M127 expansion correlated with mild COVID-19 disease outcomes. Thereby, we phenotypically and molecularly characterized a human monocyte subset marked by CD127 that retained anti-inflammatory properties within the pro-inflammatory environments, uncovering remarkable functional diversity among monocytes and signifying M127 as a potential therapeutic target for human inflammatory disorders.

## Main Text

Human monocytes and macrophages are considered major mediators of inflammation in a plethora of disease settings including infectious diseases such as COVID-19 and chronic inflammatory diseases such as rheumatoid arthritis (RA)^1–3^. During the pathological processes, inflammatory monocytes from peripheral blood origin accumulate at the sites of infection and/or inflammation and produce large quantities of pro-inflammatory mediators including cytokines and chemokines^4^, exacerbating disease outcomes by promoting the vicious inflammation cycle^5,6^. Under homeostasis, human monocyte heterogeneity has been conventionally defined by bimodal expression of CD14 and CD16^7,8^. However, understanding of functional heterogeneity of human monocytes under inflammatory conditions is limited, which imposes conceptual and technical barriers of therapeutically targeting human inflammatory diseases^9^, a particularly prominent issue amid the global menace of COVID-19 ^10^. Here, through multiomics analyses of human samples including extensive profiling at the single cell level, we defined a subset of human inflammatory monocytes uniquely marked by the expression of CD127 (thus termed M127) that were abundantly present in inflamed tissues of COVID-19 and RA patients. Mechanistic investigations and integrative computational approaches further revealed common molecular and functional features of M127 across multiple inflammatory disease conditions.

CD127, encoded by *IL7R*, is generally considered a lymphoid lineage marker that is predominantly expressed and functional on T cells and innate lymphoid cells^11^. Immunohistochemical analyses of pulmonary autopsy samples revealed minimal CD127 expression in alveoli from individuals who deceased due to non-infectious causes yet robust staining in patients succumbing to SARS-CoV-2 infection (Fig. 1a). Unexpectedly, in SARS-CoV-2 infected lung tissues, CD127 signals appeared in the regions of CD68 positivity (Fig. 1a), implying plausible expression of CD127 in monocytes/macrophages, which was confirmed by co-localization of CD127 and CD68 signals on immunofluorescently stained sections of autopsied COVID-19 lung tissues (Fig. 1b). Monocytes/macrophage (CD14^high^ CD68^high^) expression of CD127 was further corroborated at the transcriptome level by single cell RNA sequencing (scRNA-seq) analyses of bronchoalveolar lavage fluid (BALF) from nine COVID-19 patients with clinical manifestations ranging from mild (n = 3) to severe (n = 6) (Extended Data Fig. 1a). Strikingly, a distinct *IL7R*^+^ population (Fig. 1c,d) was revealed to constitute 21% of BALF monocytes/macrophages (Fig. 1e). Moreover, in contrast to the predicted dominance by lymphoid cells, the majority (64%) of *IL7R*^+^ cells in COVID-19 BALF were of monocyte/macrophage lineage (Fig. 1f).

**Figure 1.**
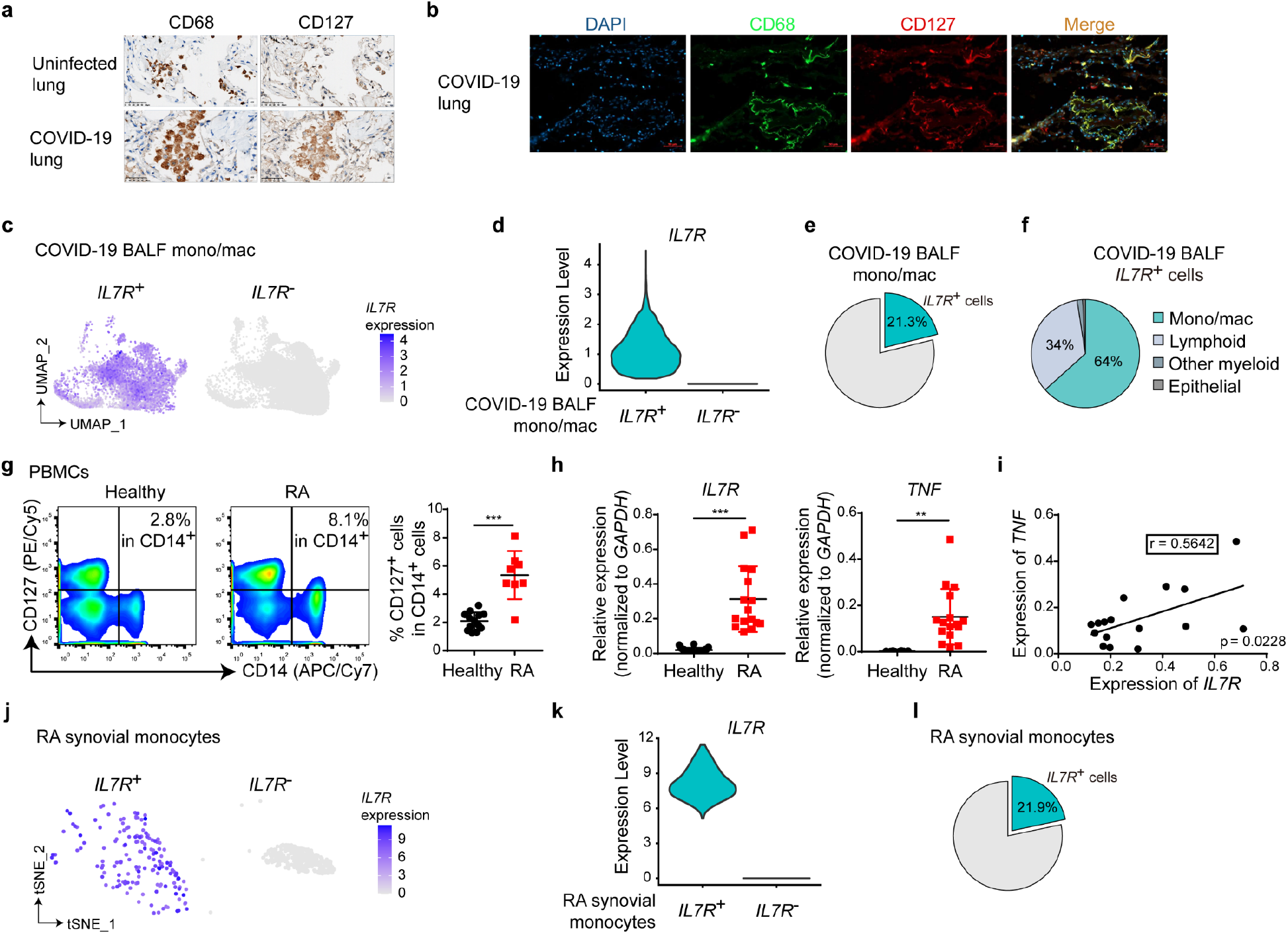
CD127high monocytes/macrophages are present in human inflammatory conditions. (**a**) Immunohistochemical analysis of CD68 and CD127 expression in lung tissues sections. Uninfected lung tissues and COVID-19 lung tissues were obtained during autopsy following the protocol described in Methods. One representative result from tissue sections of three COVID-19 cases is shown. Scale bars represent 50 μm. (**b**) Immunofluorescence staining for DAPI (blue), CD68 (green) and CD127 (red) in sections from COVID-19 lung tissues. One representative result from tissue sections of three COVID-19 cases is shown. Scale bars represent 50 μm. (**c**) UMAP projection of *IL7R*^+^ and *IL7R*^−^ monocytes/macrophages (mono/mac) in broncho-alveolar lavage fluid (BALF) from COVID-19 patients. *IL7R* expression among cells was shown by color as indicated. (**d**) Violin plot shows the expression of *IL7R* in *IL7R*^+^ and *IL7R*^−^ BALF mono/mac from COVID-19 patients. (**e**) Pie graph shows the percentage of *IL7R*^+^ cells in BALF mono/mac from COVID-19 patients. (**f**) Pie graph shows the percentages of each cell type in total *IL7R*^+^ BALF cells from COVID-19 patients. (**g**) PBMCs were isolated from the peripheral blood of healthy donors (n = 13) and rheumatoid arthritis (RA) patients (n = 8), and CD127 expression was measured by flow cytometry analysis (FACS). Representative FACS plot (left) and cumulative percentages (right) of CD127^+^ population are shown. \*\*\**P*<0.001 by unpaired Student’s *t* test. Error bars indicate means ± SD. (**h**, **i**) CD14^+^ monocytes were isolated from PBMCs of healthy donors and RA patients, and mRNA of *IL7R* and *TNF* was measured using quantitative PCR (qPCR) (**h**). Relative expression was normalized to internal control (*GAPDH*). Linear regression analysis was used to compare the expression of *TNF* and *IL7R* in CD14^+^ monocytes from RA patients (**i**). Correlation coefficient (r) and p-value for coefficient are shown in the panel. \*\**P*<0.01, \*\*\**P*<0.001 by unpaired Student’s *t* test. Error bars indicate means ± SD. (*IL7R*, Healthy n = 20 and RA n = 16; *TNF*, Healthy n = 8 and RA n = 16) (**j**) t-SNE projection of *IL7R*^+^ and *IL7R*^−^ cells in RA synovial monocytes. *IL7R* expression among cells was shown by the indicated color. (**k**) Violin plot shows the expression of *IL7R* in *IL7R*^+^ and *IL7R*^−^ RA synovial monocytes. (**l**) Pie graph shows the percentage of *IL7R*^+^ cells in RA synovial monocytes.

To validate whether CD127 expression on monocytes/macrophages can be generalized to other human inflammatory conditions such as RA, peripheral blood mononuclear cells (PBMCs) from 16 anti-inflammatory treatment naïve RA patients were analyzed for CD127 expression. Compared with the minimal levels of CD127 in CD14^+^ monocytes from healthy donors, all RA patients examined displayed markedly elevated expression of CD127 on blood monocytes at the protein and mRNA levels (Fig. 1g,h). In RA blood monocytes, *IL7R* expression correlated with the expression of a major pathogenic factor, TNF (Fig. 1h,i) ^12^. To probe CD127 at the primary sites of inflammation, RA synovial tissue scRNA-seq data sets from 18 patients^13^ (Extended Data Fig. 1b) were analyzed for *IL7R* expression, which showed strong positivity in a distinct population (Fig. 1j,k) that represented nearly 22% of synovial monocytes (Fig. 1l). Taken together, expression of CD127 on a subset of inflammatory monocytes/macrophages is likely a hallmark of human inflammatory conditions testified in multiple disease settings (COVID-19 and RA) and multiple tissues (infected lungs, peripheral blood and inflamed joints).

In order to pursue in-depth investigation of CD127^+^ monocytes/macrophages, we wished to recapitulate such phenotypes *in vitro* using infectious and inflammatory stimuli. Stimulation of human PBMCs from healthy donors with toll-like receptor (TLR) ligands led to drastic increase of monocytic CD127 at the protein and mRNA levels, with the exception of TLR3 agonist poly(I:C) (Fig. 2a,b). Upregulation of CD127 was dynamic, peaking around 6 h post LPS stimulation (Fig. 2c,d), dependent on canonical TLR signaling modules such as IKK and p38 (Extended Data Fig. 2a-e), and observed in all three currently defined human monocyte subpopulations (Fig. 2e and Extended Data Fig. 2f,g). In contrast to human monocytes, LPS failed to upregulate CD127 in murine peripheral blood monocytes (Extended Data Fig. 3a-c) and macrophages from multiple tissue sources (data not shown), suggesting human-specific nature of CD127 induction. In addition to TLR stimulations that mimicked infectious conditions, we wished to identify factors that led to CD127 expression in chronic inflammatory diseases and pursued TNF as a plausible candidate as its levels correlated with CD127 expression (Fig. 1i). TNF treatment consistently upregulated CD127 in monocytes, albeit to a lesser extent than LPS (Fig. 2f,g). Importantly, clinically applied TNF blockade treatment significantly reduced *IL7R* expression in RA monocytes (Fig. 2h), further solidifying a role for TNF in upregulation of CD127 *in vivo*. Next, we wished to investigate whether CD127 in activated monocytes was functional given that another subunit of IL-7 receptor, the *IL2RG*-encoding common gamma chain, was constitutively expressed in human monocytes (Extended Data Fig. 3d). IL-7 treatment robustly induced STAT5 tyrosine phosphorylation in LPS-activated monocytes but not in resting monocytes (Fig. 2i) with T cells serving as positive controls (Extended Data Fig. 3e), indicating that activated human monocytes were competent for IL-7 receptor signaling.

**Figure 2.**
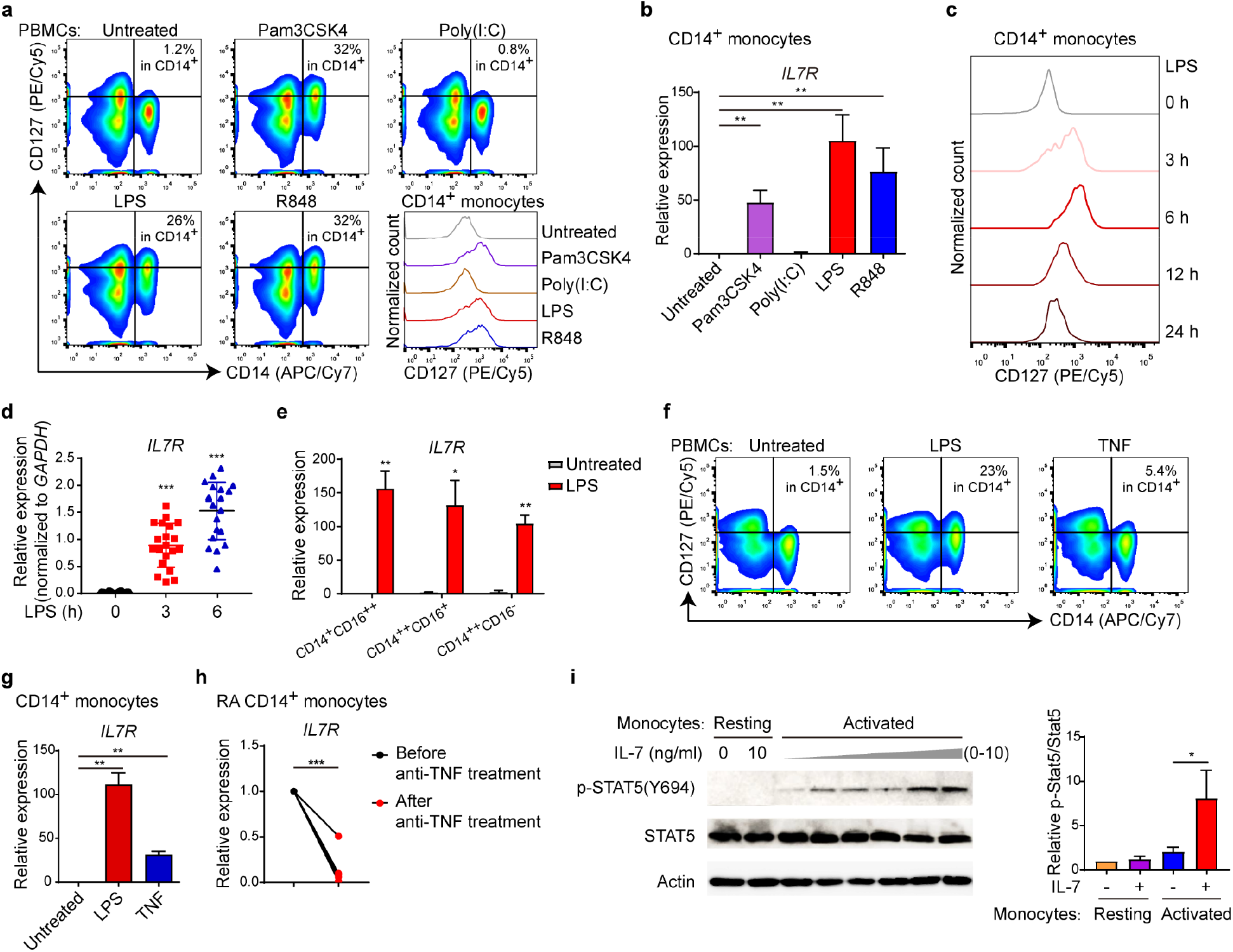
CD127high monocytes are inducible by inflammatory stimuli and competent for CD127-STAT5 signaling. (**a**) PBMCs from healthy blood donors were stimulated with Pam3CSK4 (100 ng/ml), Poly(I:C) (1 μg/ml), LPS (10 ng/ml) or R848 (1 μg/ml) for 6 h, and CD127 expression was measured by FACS. Histograms in bottom right shows the CD127 staining signal in CD14^+^ cells under each condition as indicated. One representative FACS result from three biological replicates is shown. (**b**) CD14^+^ monocytes from healthy donors’ PBMCs were stimulated with Pam3CSK4 (100 ng/ml), Poly(I:C) (1 μg/ml), LPS (10 ng/ml) or R848 (1 μg/ml) for 3 h. The mRNA of *IL7R* was measured by qPCR. Relative expression was normalized to internal control (*GAPDH*) and expressed relative to untreated sample. \*\**P*<0.01 by paired Student’s *t* test. Data are shown as means ± SD of four independent experiments with one healthy donor for each data set. (**c**) CD14^+^ monocytes from healthy donor’s PBMCs were treated with 10 ng/ml LPS for various time points as indicated, and the expression of CD127 was measured by FACS. (**d**) CD14^+^ monocytes were treated with 10 ng/ml LPS for 3 h and 6 h, the mRNA of *IL7R* was measured by real time qPCR. Relative expression was normalized to *GAPDH* as internal control. Each data point represented an independent experiment from one healthy donor. \*\*\**P*<0.001 by unpaired Student’s *t* test. Error bars indicate means ± SD of 20 independent experiments. (**e**) Three different monocyte subsets were FACS-sorted from PBMCs as shown in Extended Data Fig. 2f. The sorted cells were treated with or without 10 ng/ml LPS for 3 h. The expression of *IL7R* was measured by q-PCR. Relative expression was normalized to internal control (*GAPDH*) and expressed relative to LPS-untreated CD14^+^CD16^++^ sample. \*\**P*<0.01, \**P*<0.05 by unpaired Student’s *t* test. Data are shown as means ± SD of three independent experiments with one healthy donor for each data set. (**f**, **g**) PBMCs from healthy donor were treated with LPS (10 ng/ml) or recombinant human TNF (100 ng/ml). Upon 6 h LPS or TNF treatment, CD127 expression was measured by FACS. Representative FACS distribution is shown (**f**). mRNA of *IL7R* was measured in 3 h LPS or TNF treated monocytes by qPCR (**g**). Relative expression was normalized to internal control (*GAPDH*) and expressed relative to untreated sample. \*\**P*<0.01 by paired Student’s *t* test. Data are shown as means ± SD of three independent experiments with one healthy donor for each data set. (**h**) mRNA level of *IL7R* was measured by real time qPCR in monocytes from RA patients (n = 5) before and after Etanercept anti-TNF treatment for 2 months. Relative expression was normalized to internal control (*GAPDH*). \*\*\**P*<0.001 by paired Student’s *t* test. (**i**) CD14^+^ monocytes were pre-treated with or without LPS for 6 h, then followed by various doses of recombinant human IL-7 (from 1 pg/ml to 10 ng/ml) for 30 min. STAT5 activation was detected by western blotting. Actin was used as loading control. One representative experiment out of three biological replicates is shown (left). The protein level of p-STAT5(Y694) was quantified by densitometry, normalized to total STAT5 protein and expressed relative to untreated (without LPS and IL-7) sample (right). \**P*<0.05 by paired Student’s *t* test. Data were expressed as mean ±SD of three independent experiments with one healthy donor for each data set.

**Figure 3.**
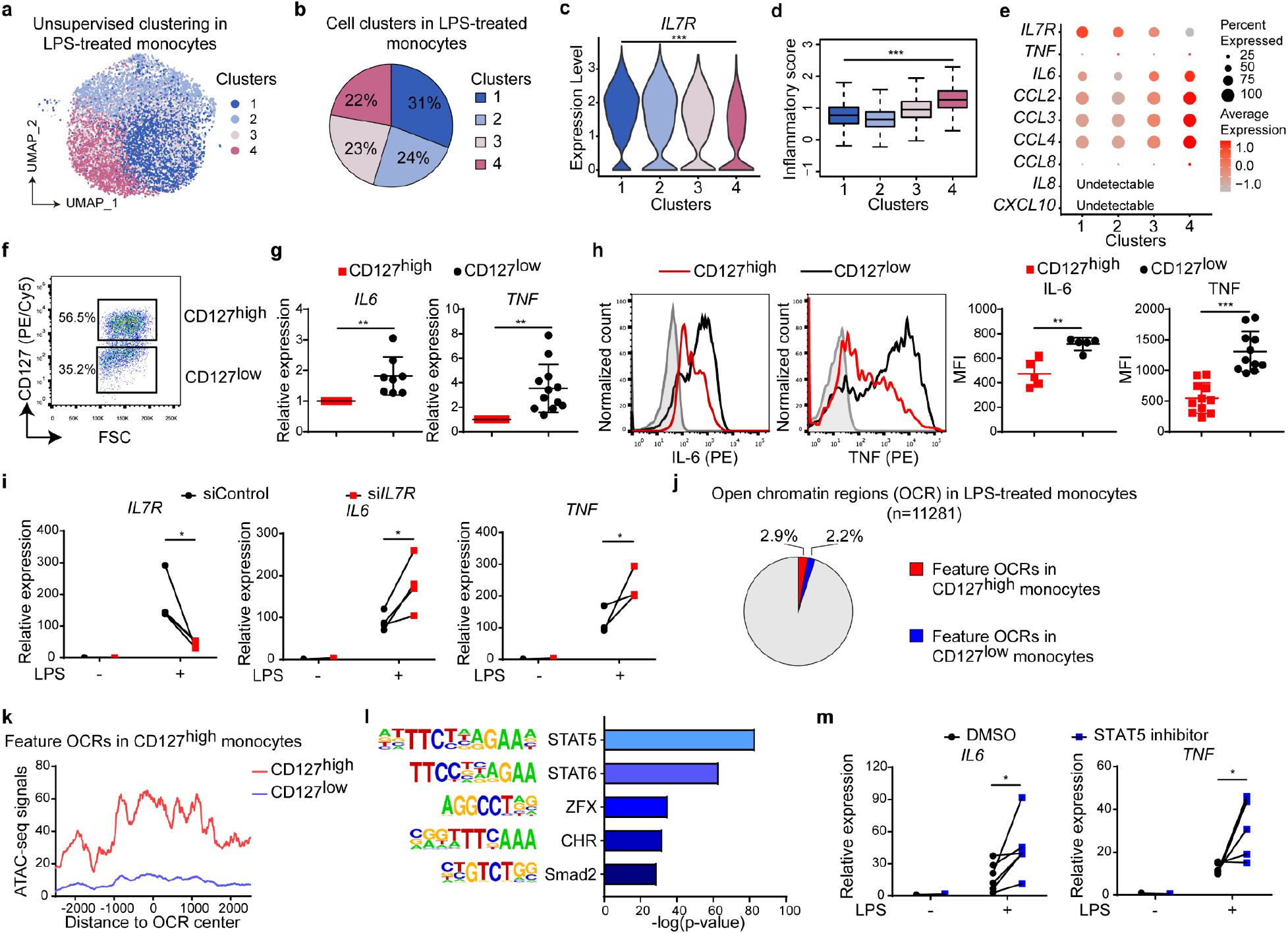
CD127 imposes heterogeneity in monocyte inflammatory responses mediated by CD127-STAT5 axis. (**a**) UMAP projection of LPS 6 h treated human CD14^+^ monocytes that were subgrouped by unsupervised cluster analysis of scRNA-seq data. (**b**) Pie graph shows the percentages of each monocyte cluster shown in in LPS-treated monocytes. (**c**) Violin plot shows the expression of *IL7R* among four clusters of LPS-treated monocytes. \*\*\**P*<0.001 by Wilcoxon rank-sum test. (**d**) Inflammatory score was defined by average expression of eight inflammatory genes, and box plot shows the inflammatory score distribution among four clusters of LPS-treated monocytes. \*\*\**P*<0.001 by Wilcoxon rank-sum test. (**e**) Heat map shows the expression of *IL7R* and eight inflammatory genes for inflammatory score calculation among four clusters of LPS-treated monocytes. The scaled average expression levels for each gene and percentages of cells expressing each gene in each cluster were represented by color and size of the corresponding dots, respectively. (**f**, **g**) CD127^high^ and CD127^low^ populations were isolated from 6 h LPS-stimulated CD14^+^ monocytes by FACS (**f**) and the mRNA levels of *IL6* and *TNF* in two populations were measured by qPCR (**g**). Relative expression was normalized to internal control (*GAPDH*) and shown relative to CD127^high^ sample. \*\**P*<0.01, \*\*\**P*<0.001 by paired Student’s *t* test. Data are shown as means ± SD of eight independent experiments for *IL6* and twelve independent experiments for *TNF*. (**h**) Protein levels of IL-6 and TNF in CD127^high^ and CD127^low^ populations were measured by GolgiStop-utilized intracellular staining and analyzed by flow cytometry. One representative FACS plot (left) and cumulative mean fluorescence intensities (MFI) from five independent experiments for IL-6 and eleven independent experiments for TNF (right) are shown. \*\**P*<0.01, \*\*\**P*<0.001 by paired Student’s *t* test. Data are shown as means ± SD. (**i**) CD14^+^ monocytes were transfected with negative control or *IL7R* specific short interfering RNA (siControl or si*IL7R*). Two days post transfection, cells were stimulated with LPS (10 ng/ml) for 3 h, and the mRNA of *IL7R*, *IL6* and *TNF* were measured by using real time qPCR. Relative expression was normalized to internal control (*GAPDH*) and expressed relative to LPS untreated siControl sample. \**P*<0.05 by paired Student’s *t* test. (**j**) Three healthy donors’ CD127^high^ and C127^low^ monocytes were sorted as in (F), and generation of ATAC-seq data sets were performed for two monocyte populations from each donor. Pie graph shows the percentages of feature open chromatin regions (OCR) in CD127^high^ and C127^low^ monocytes in total OCRs in LPS-treated monocytes by statistical analyses for three ATAC-seq data sets. (**k**) Three ATAC-seq data sets were combined, and average ATAC-seq signals were calculated in combined ATAC-seq data set for CD127^high^ and C127^low^ monocytes around feature OCRs in CD127^high^ monocytes. (**l**) Motif enrichment analysis in CD127^high^ monocyte featured open chromatin regions. Top 5 most enriched transcription factor binding motifs are shown, and x-axis is −log10(p-value) for each enriched motif. Binomial distribution was used for p-value calculation. (**m**) CD14+ monocytes were pretreated with STAT5 inhibitor (100 μM) for 2 h and then were stimulated with LPS (10 ng/ml). mRNA levels of *IL6* (LPS 6 h) and *TNF* (LPS 3 h) were measured using qPCR. \**P*<0.05 by paired Student’s t test. The results from six independent experiments are shown.

Given the heterogeneity of CD127 expression, we reasoned that activated human monocytes may display functional diversity and subjected LPS-activated monocytes to single cell expression profiling. Unsupervised hierarchical clustering revealed four groups of cells with differential gene expression patterns (Fig. 3a,b). Interestingly, *IL7R* exhibited a gradient pattern among four clusters, with the highest expression in cluster 1 and the lowest in cluster 4 (Fig. 3c). To quantitatively assess the inflammatory phenotypes of these cells, we devised a numeric index ‘inflammatory score’ based on an algorithm^14^ reflecting expression levels of 8 representative prototypical inflammatory genes (see ‘Methods’ for details). Inflammatory score inversely correlated with the expression of *IL7R*, with the highest level shown for cluster 4 that exhibited the lowest level of *IL7R* (Fig. 3d), a trend that could also be clearly visualized for individual inflammatory genes (Fig. 3e). To validate the differences of inflammatory responses observed from single cell analyses, we sorted CD127^high^ and CD127^low^ LPS-activated monocytes (Fig. 3f). Consistent with the single cell results, CD127(*IL7R*)^low^ monocytes produced significantly higher levels of inflammatory mediators such as IL-6 and TNF (Fig. 3g,h), demonstrating that within a highly defined system consisting of purified monocytes and a single stimulus, human monocytes displayed the remarkably diverse range of inflammatory responses.

Having observed the inverse correlation between CD127 and inflammatory responses, we wished to examine whether CD127 was causally related to inflammation by knocking down *IL7R* in monocytes with RNA interference. Downregulation of *IL7R* resulted in markedly upregulated expression of *IL6* and *TNF* (Fig. 3i), implicating a negative role for monocytic CD127 in inflammation. To characterize the chromatin accessibility landscape of CD127^high^ and CD127^low^ monocytes, we subjected two populations to ATAC-seq (Extended Data Fig. 4a) and identified a highly specified fraction of open chromatin regions in CD127^high^ monocytes (Fig. 3j,k) that displayed enhancer-like features marked by H3K27ac and H3K3me1 modifications^15^ (Extended Data Fig. 4b,c) and enriched in binding motifs for several transcription factors with STAT5 being the most prominent (Fig. 3l). Pharmacological inhibition of STAT5 led to upregulation of *IL6* and *TNF* expression (Fig. 3m), in line with the effects of *IL7R* abrogation. To elucidate the mechanisms underlying CD127-STAT5-mediated effects, CD127^high^ and CD127^low^ population were profiled by RNA-seq (Extended Data Fig. 5a,b), revealing that transcription factor c-Maf, encoded by the *MAF* gene, was relatively highly expressed in CD127^high^ cells in a STAT5-dependent manner (Extended Data Fig. 5c,d). Knocking down *MAF* expression resulted in upregulation of *IL6* and *TNF* (Extended Data Fig. 5e,f). Of note, *MAF* is a direct STAT5 target gene as shown by occupancy of STAT5 at a consensus binding site upstream of the *MAF* transcription start site (Extended Data Fig. 5g). Together, the above results implicated that the CD127-STAT5-c-Maf axis exerted anti-inflammatory effects, contributing to the functional heterogeneity in human inflammatory monocytes.

**Figure 4.**
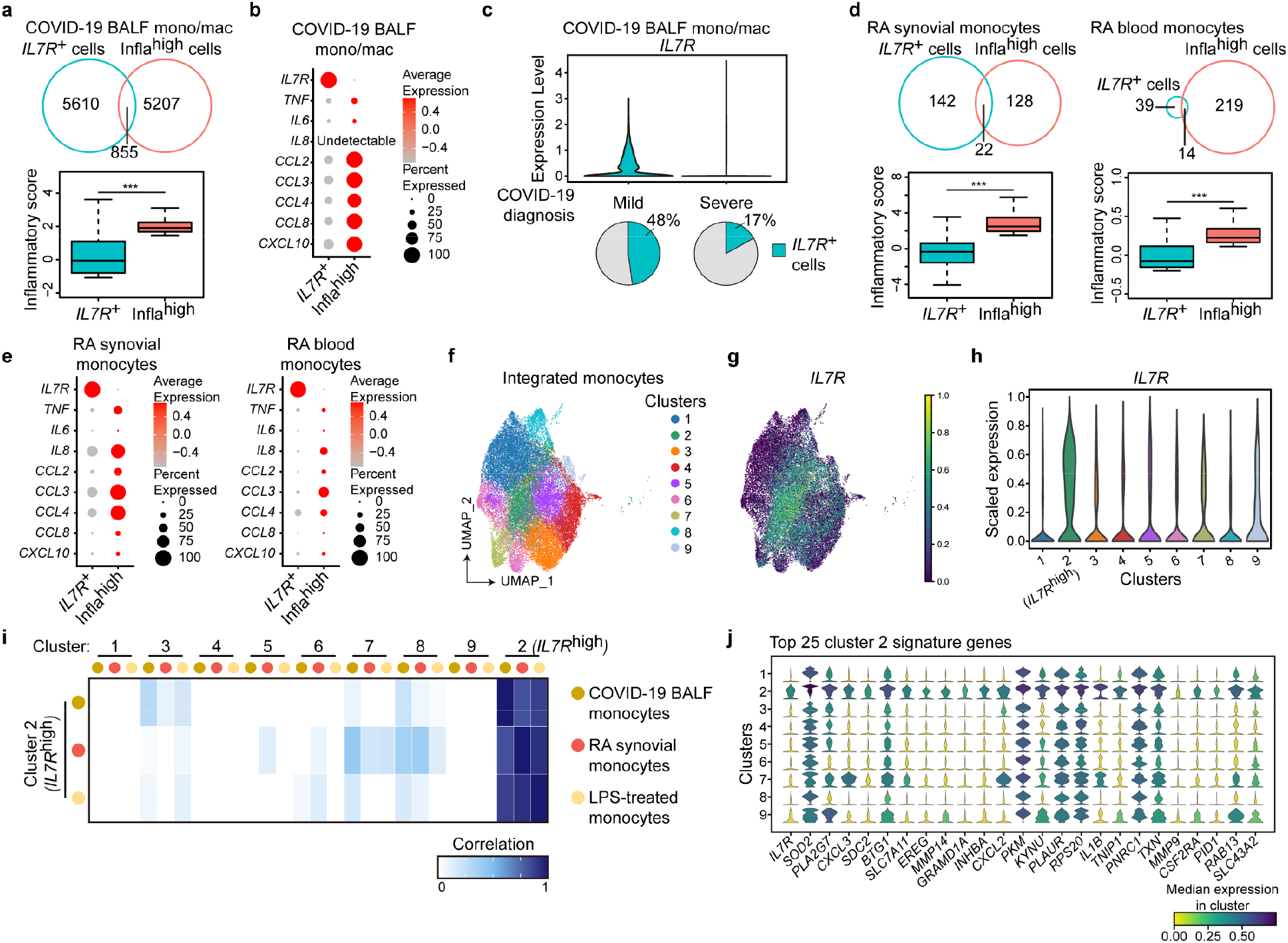
CD127high monocytes are anti-inflammatory in human diseases. (**a**, **d**) Infla^high^ cells for each disease condition were subgrouped as top 20% inflammatory cells by inflammatory score in mono/mac from COVID-19 BALF, RA synovial monocytes and RA blood monocytes, respectively. Venn diagrams (upper) show the extent of overlap between *IL7R*^+^ cells and infla^high^ cells, and box plots (bottom) show the inflammatory score distribution in *IL7R*^+^ cells and infla^high^ cells in mono/mac from COVID-19 BALF (**a**), RA synovial monocytes and RA blood monocytes (**d**) as indicated. \*\*\**P*<0.001 by Wilcoxon rank-sum test. (**b**, **e**) Heat maps show the expression of *IL7R* and eight inflammatory genes for inflammatory score calculation in *IL7R*^+^ cells and infla^high^ cells in mono/mac from COVID-19 BALF (B), RA synovial monocytes and RA blood monocytes (**e**) as indicated. The scaled average expression levels for each gene and percentages of cell expressing each gene in each group were represented by color and size of the corresponding dots, respectively. (**c**) Mono/mac cells from COVID-19 patients’ BALF were subgrouped by the diagnosed disease severity. Violin plot (upper) shows the expression of *IL7R* in BALF mono/mac from mild or severe COVID-19 patients. Pie graphs on the bottom show the percentage of *IL7R*^+^ cells in BALF mono/mac from mild or severe COVID-19 patients as indicated. (**f**) UMAP projection of integrated monocytes from COVID-19 BALF, RA synovial cavity and LPS-treated monocytes. Nine monocyte clusters by unsupervised clustering were indicated by different colors in plot. (**g**) UMAP projection of integrated monocytes in (**f**). *IL7R* expression among cells were expressed by color as indicated. (**h**) Violin plot shows the expression of *IL7R* among nine clusters of integrated-monocytes in (**f**). Cluster 2 were named as *IL7R*^high^ cluster by highest *IL7R* expression among clusters. (**i**) For populations of COVID-19 BALF monocytes, RA synovial monocytes and LPS-treated monocytes, heat map shows the correlation between cluster 2 cells in each monocytes population and all nine clusters in each monocytes population, respectively. (**j**) The stacked violin plot shows the expression of top 25 cluster 2 signature genes in all nine monocyte clusters in (**f**). Median expression levels for each gene in each cluster were expressed by color as indicated.

Upon characterizing the hypo-inflammatory phenotypes of CD127^high^ cells *in vitro*, we next wished to validate these findings *in vivo* in human disease settings. In BALF monocytes/macrophages from COVID-19 patients, *IL7R*^+^ population did not extensively overlap with the highly inflammatory cells and exhibited minimal inflammatory properties (Fig. 4a,b). Consistent with the *in vitro* observations, *IL7R*^+^ cells expressed heightened levels of *MAF* relative to the highly inflammatory monocytes (Extended Data Fig. 6a). Importantly, in mild COVID-19 patients, *IL7R*^+^ cells constituted 48% of all BALF monocytes/macrophages yet in stark contrast, such percentage was merely 17% in severe COVID-19 patients (Fig. 4c). The differential *IL7R* patterns in monocytes were not due to the global differences in expression levels as lymphocytic *IL7R* did not significantly differ between mild and severe patients (Extended Data Fig. 6b). These results indicated that CD127 indeed marked a population of hypo-inflammatory monocytes/macrophages *in vivo* and suggested that the prevalence of *IL7R*^+^ monocytes/macrophages likely correlated with subdued inflammation and favorable disease outcomes. In addition to COVID-19, *IL7R*^+^ population in RA synovial monocytes and peripheral blood monocytes were also largely non-overlapping with the highly inflammatory counterparts and displayed minimal inflammatory features (Fig. 4d,e and Extended Data Fig. 6c,d).

To extrapolate common features of *IL7R*^+^ monocytes/macrophages from various disease conditions, a recently developed bioinformatics method specializing in incorporating single cell sequencing data sets from multiple sources^16^ was used to run integrated analyses of three data sets: COVID-19 BALF monocytes/macrophages, RA synovial monocytes and *in vitro* LPS-activated monocytes. Unsupervised clustering identified 10 cellular subsets, with the cluster 10 classified as tissue resident alveolar macrophages from the COVID-19 samples (Extended Data Fig. 6e-g) and thus being excluded from the subsequent analyses intending to discover common monocyte characteristics regardless of tissue origins. The remaining 9 clusters (1-9) represented integrated monocyte subsets present in all three tissue sources (Fig. 4f), with cluster 2 prominently featured by high *IL7R* expression (Fig. 4g,h) and sharing high degree of similarity among three conditions (Fig. 4i). Common signature genes of cluster 2 revealed a profile that was distinct from any of the currently known monocyte/macrophage subsets, with *IL7R* unambiguously identified as the top marker gene (Fig. 4j). Of note, cluster 2 phenotypically differed from the ‘M2-like’ cells defined by markers such as CD163 (Extended Data Fig. 6h).

In summary, we identified a subset of monocytes marked by CD127 in human infectious and inflammatory diseases but not in mice, and named this subset M127 (Extended Data Fig. 7). We further characterized the inflammatory signals that induced M127 and recapitulated M127 phenotypes with an *in vitro* system. CD127 did not only serve as a surface marker for this population but also actively transmitted local IL-7 cues^17^ to promote a STAT5-coordiated anti-inflammatory program, resulting in hypo-inflammatory phenotypes amid the overall inflammatory tissue environments. As of the knowledge of the current study, M127 from multiple disease conditions and multiple tissues such as COVID-19 lungs and RA joints shared common functional features and gene signatures, albeit it would be interesting and desirable to assess whether the depicted M127 phenotypes could be observed in additional human disease settings. Given the unique presence of this population in human inflammatory diseases, especially the correlation of M127 expansion with favorable disease outcomes in COVID-19, it is highly conceivable to propose M127 as a potential therapeutic target for inflammatory disorders.

## Methods

### Cell culture and reagents

PBMCs of anonymous healthy donors were isolated from buffy coats purchased from the Beijing Red Cross Blood Center using density gradient cell separation by Ficoll (Lymphoprep^™^, STEMCELL Technologies) following the protocol approved by the Institutional Review Board of School of Medicine, Tsinghua University. The private information of anonymous blood donors was inaccessible to investigators. PBMCs of RA patients were obtained from Peking Union Medical College Hospital using the protocol that was approved by the Institutional Review Board of Peking Union Medical College Hospital. CD14^+^ Monocytes were further isolated from PBMCs using anti-CD14 magnetic beads (130-050-201, Miltenyi Biotec). CD14^+^ monocytes were cultured in RPMI 1640 medium (10040CM, Corning) supplemented with 10% (vol/vol) fetal bovine serum (FBS) (Gibco) and human recombinant M-CSF (300-25, Peprotech) (10 ng/ml). LPS (*Escherichia coli* O127:B8, Sigma-Aldrich), human recombinant IL-7 (200-07, Peprotech), human recombinant TNF (H8916, Sigma-Aldrich) or chemical inhibitors (SB203580 from Selleck, STAT5 Inhibitor from Santa Cruz and Bay 11-7082 from Sigma-Aldrich) were used as indicated for various experiments.

### Collection of lung tissues and immunohistochemistry

Two cases of uninfected lung tissues and 3 cases of COVID-19 lung tissues were from Biobank of Southwest Hospital, Third Military Medical University (Army Medical University). COVID-19 lung tissues were obtained during autopsy of the patients succumbing to SARS-CoV-2 infection. Pathologically normal lung tissues from pulmonary bulla patients were used as uninfected controls. Tissue collection and the following histological analyses were approved by the ethics committee of Southwest Hospital, Third Military Medical University (Army Medical University), and were in accordance with regulations issued by the National Health Commission of China and the Helsinki Declaration. Lung tissue sections were stained with hematoxylin for assessment of pulmonary architecture, and anti-CD68 (ab201340, abcam) and anti-CD127 (PA5-97870, Invitrogen) antibodies were used for immunohistochemistry. Specifically, lung sections were deparaffinized and rehydrated. Antigen retrieval was performed with the Improved Citrate Antigen Retrieval Solution (Beyotime) and incubated with H_2_O_2_ in dark for 15 min to block endogenous peroxidase activity. Slides were blocked with 10% goat serum in TBS for 30 min at room temperature and stained with primary antibodies overnight at 4°C. Slides were washed three times with 0.1% TBS-Tween before incubation with HRP-conjugated secondary antibodies. Stained slides were washed again in PBS and stained with DAB (TIANGEN) in conjunction with a hematoxylin counterstain (Solarbio). After dehydration, sections were mounted in neutral balsam.

### Immunofluorescence histology

COVID-19 lung tissues were collected as described above, and were washed and fixed overnight at 4°C in a solution of 1% paraformadehyde in PBS. The tissues were incubated in a solution of 30% sucrose in PBS and the mixture of 30% sucrose and OCT compound 4583 (Sakura Finetek) separately at 4°C overnight. The samples were then embedded in OCT, frozen in a bath of ethanol cooled with liquid nitrogen and stocked at −80°C. Frozen samples were cut at 10-μm thickness and collected onto slides. Slides were dried at 50°C for 30 min and fixed in 1% paraformaldehyde for 10 min and processed for staining. The tissues were permeabilized in PBS/0.3% Triton X-100/0.3 M glycine at 37°C for 30 min and blocked in PBS/5% goat serum at room temperature for 1 h. The tissues were then incubated with indicated primary antibodies diluted (anti-CD68, 1:100; anti-CD127, 1:500) in PBS/5% goat serum at 4°C overnight, and washed in PBS/0.2% Tween-20 at room temperature for 30 min three times. The tissues were incubated with Alexa dye-conjugated secondary antibodies (Alexa Fluor^™^ 488 goat anti-mouse IgG, 1:500, B40941, Life Technologies; Alexa Fluor^™^ 555 goat anti-rabbit IgG, 1:500, A27039, Invitrogen) and DAPI (1:200, C0060-1, Solarbio) in PBS/0.5% BSA at room temperature for 2 h and washed in PBS/0.2% Tween-20 at room temperature for 1 h five times before mounting with SlowFade Diamond Antifade Mountant (S36963, Life Technologies).

### Bronchoalveolar lavage fluid (BALF) collection for single cell RNA sequencing (scRNA-seq)

Nine COVID-19 patients were enrolled from the Shenzhen Third People’s Hospital. BALF collection from COVID-19 patients and healthy donors and following studies were conducted according to the principles expressed in the Declaration of Helsinki. Ethical approval was obtained from the Research Ethics Committee of Shenzhen Third People’s Hospital (2020-112). Diagnosis of COVID-19 was based on clinical symptoms, exposure history, chest radiography and SARS-CoV-2 RNA positivity. Disease severity was defined as moderate, severe and critical, according to the ‘Diagnosis and Treatment Protocol of COVID-19’ by the National Health Commission of China. Approximately 20 ml of BALF was obtained for each patient. BALF was directly processed within 2 h and all operations were performed in a BSL-3 laboratory. BALF cells were collected, counted, re-suspended, and subsequently processed for scRNA-seq library construction as described in our previous study^18^. According to the clinical diagnosis, nine enrolled patients included three moderate cases, one severe case and five critical cases. For the subsequent analyses of scRNA-seq data, given that there was only one clinically defined severe case, the patients were stratified into mild (n = 3) and severe (n = 6 including both severe and critical cases) groups.

### Mice

The laboratory animal facility at Tsinghua University has been accredited by AAALAC (Association for Assessment and Accreditation of Laboratory Animal Care International), and the IACUC (Institutional Animal Care and Use Committee) of Tsinghua University approved the protocol used in this study for blood collection from mouse cheeks. C57BL/6J mice were bred and housed in isolated ventilated cages (maxima six mice per cage) at the specific pathogen free facility at Tsinghua University. The mice were maintained on a 12/12-h light/dark cycle, 22–26 °C, 40–70% humidity with sterile pellet food and water ad libitum.

### RNA extraction and quantitative PCR

Total RNA was extracted from cells using TRIzol^®^ Reagent according to the manufacturer’s procedure, and total RNA was reverse-transcribed to cDNA with Moloney Murine Leukemia Virus Reverse Transcriptase (2641B, TAKARA). Real-time quantitative PCR (qPCR) was performed in duplicates with SYBR Green Master Mix (A25742, Applied Biosystems) on StepOnePlus thermal cycler (Applied Biosystems). Primer sequences are listed in the Extended Data Table 1.

### Flow cytometry

Upon indicated treatment, cells were collected and washed with staining buffer (PBS with 0.5% BSA and 2 mM EDTA). Then, the surface markers were stained with the indicated fluorochrome-conjugated antibodies in 1:400 dilution for 30 min on ice in the dark. After staining, cells were washed three times with staining buffer and re-suspended in PBS for analysis in BD FACSFortessa or for fluorescence-activated cell sorting (FACS) in BD FACSAria III. Further data analysis was implemented using Flowjo software (Tree star). For intracellular staining, cells were treated with Golgistop (554724, BD Biosciences) for 4–5 h before collection. The routine stainings for surface markers were performed, after which the cells were fixed with 100 μl/tube Fixation Buffer (420801, Biolegend) for 25 minutes at room temperature, and the fixed cells were permeabilized and stained in 1 × Permeabilization Wash Buffer with fluorochrome-conjugated antibodies for 30 minutes on ice in the dark. The fixed and intracellularly stained cells were washed twice with 1 × Permeabilization Wash Buffer and suspended in PBS for analysis. The fluorochrome-conjugated antibodies for targets of interest and fluorochrome-conjugated isotype control antibodies are listed in Extended Data Table 2.

### Immunoblotting

Whole-cell lysates were prepared by direct lysis in sodium dodecyl sulfate (SDS) loading buffer. All samples for immunoblotting were denatured at 95 °C for 10 min. For immunoblotting analysis, denatured cell lysates were separated by 10% SDS polyacrylamide gel electrophoresis and transferred to a polyvinylidene fluoride membrane (Millipore) for probing with specific primary antibodies and HRP-conjugated secondary antibodies. SuperSignal™ West Pico Chemiluminescent Substrate (34580, Thermo Fisher Scientific) was used for detection. Relative density of blotting bands was quantified using Image J (v1.52a). Antibodies used for probing proteins of interest are listed in Extended Data Table 2.

### RNA interference

Immediately after isolation, primary human monocytes were nucleofected with On-Target plus SMARTpool siRNA purchased from Dharmacon Inc. specific for *IL7R*, *MAF* or *MAP3K3*. Non-targeting siRNA from GenePharma was used as control. Human Monocyte Nucleofector buffer (V4XP-3024, Lonza) and the Lonza 4D-Nucleofector™ platform were used according to the manufacturers’ instructions with human monocytes nucleofection program. The nucleofected monocytes were cultured in RPMI 1640 medium (Corning) supplemented with 10% (vol/vol) FBS (Gibco) and human recombinant M-CSF (Peprotech) (20 ng/ml) for 48 h before the following experiments.

### Chromatin immunoprecipitation (ChIP) assay

For STAT5 ChIP assays, THP-1 cells were stimulated with LPS (100 ng/ml) for 6 h and subsequently with IL-7 (10 ng/ml) for 30 min. 10–20 × 10^6^ cells per condition were fixed in 1% methanol-free formaldehyde (Thermo Scientific) for 8 min at room temperature followed by quenching with 125 mM glycine for another 5 min. ChIP assay was performed using the SimpleChIP enzymatic ChIP kit (Cell Signaling Technology) according to the manufacturer’s instructions. The DNA-protein complexes were immunoprecipitated using 5.0 μl per sample of STAT5 antibody (9363S, Cell Signaling Technology), and IgG (2729P, Cell Signaling Technology) control was performed in an equally allocated DNA-protein complexes fraction as STAT5 ChIP samples. The immunoprecipitated DNA fragments were extracted with QIAquick PCR purification kit (QIAGEN) and subjected to qPCR assay for enrichment detection in *MAF* transcription start site (TSS) upstream GAS motif with primer pair, forward-AAGTGCAGTGCTATAAAGTTGTTT and reverse-ATGTTCAAGACGCTGGCTTA.

### RNA-seq

Human CD14^+^ monocytes were stimulated with LPS 10 ng/ml for 6 h, and CD127^high^ and CD127^low^ populations for each donor were sorted by FACS. Total RNA was extracted from CD127^high^ and CD127^low^ cells using TRIzol^®^ Reagent (Thermo Fisher Scientific) according to the manufacturer’s procedure, and RNA samples were processed for library construction with TruSeq mRNA-seq Sample Preparation Kit (Illunima) and sequencing in BGI Genomics Co., Ltd. on a BGISEQ-500RS platform. Three independent sets of paired samples collected from three healthy donors were subjected to RNA-seq and the subsequent bioinformatics analyses.

### Single cell RNA sequencing for RA PBMCs and LPS-activated monocytes

After the isolation or treatment, cells were frozen in FBS + 10% DMSO for preservation in liquid nitrogen. The frozen cells were processed for scRNA-seq in BGI Genomics Co., Ltd. Single-cell capturing and downstream library constructions were performed using Chromium Single Cell 3’ Reagent kits (10x Genomics) according to the manufacturer’s protocol. The constructed libraries were sequenced on a BGI MGISEQ2000 platform.

### Assay for transposase-accessible chromatin coupled with high-throughput sequencing (ATAC-seq)

Human CD14^+^ monocytes from healthy donors were stimulated with 10 ng/ml of LPS for 6 h, and CD127^high^ and CD127^low^ populations for each donor were sorted by FACS. Cells were pelleted by centrifugation for 10 min at 500 g 4°C using a swing rotor with low acceleration and brake settings. Cell pellets were washed once with 1 × PBS and cells were pelleted again by centrifugation using the previous settings. Cell pellets were re-suspended in 50 μl of lysis buffer (10 mM Tris-HCl pH 7.4, 10 mM NaCl, 3.0 mM MgCl2, 0.5% NP-40) and nuclei were pelleted by centrifugation for 30 min at 500 g, 4°C using a swing rotor with low acceleration and brake settings. Supernatant was discarded and nuclei were re-suspended in 50 μl reaction buffer containing 5.0 μl Tn5 transposase and 10 μl of 5 × TTBL buffer (TruePrepTM DNA Library Prep Kit V2 for Illumina, Vazyme Biotech). The reaction was incubated at 37°C for 30 min. After the tagmentation, the transposed DNA fragments were purified by 1 × AMPure XP beads (Beckman Coulter). PCR was performed to amplify the libraries for 9 cycles using the following PCR conditions: 72°C for 3min; 98°C for 30 s; and thermocycling at 98°C for 15 s, 60°C for 30 s and 72°C for 3 min; following by 72°C 5 min. After the PCR reaction, libraries were purified with the 0.5 × and 1.2 × AMPure XP beads. DNA concentrations were measured with StepOnePlusTM Real-Time PCR System (Life Technologies) and library sizes were determined using Agilent 2100 Bioanalyzer. Libraries were sequenced on an Illumina Hiseq X-ten platform for an average of 20 million unique reads per sample. Three independent sets of paired samples collected from three healthy donors were subjected to ATAC-seq and the subsequent bioinformatics analyses.

### Next-generation sequencing (NGS) data alignment

ATAC-seq pair-end reads were collected. Adapter sequences were trimmed from the ends of reads by Cutadapt (v1.14), and the reads that failed to pass the quality control (Q > 10) were discarded. H3K27ac and H3K4me1 ChIP-seq data sets were downloaded from NCBI GEO DataSet under the GEO accessions: GSE85245^19^. SRA files were converted to fastq files using fastq-dump included in SRA toolkit. Pair-end ATAC-seq reads were aligned to human genome (UCSC hg38) using Bowtie2 (v2.2.5)^20^ to generate alignment files of uniquely mapped pair-end fragments with maximum length in 1000 bp and no more than one mismatch for each alignment seed with 15 bp in length. ChIP-seq reads in fastq files were aligned to human genome (UCSC hg38) using Bowtie (v1.1.2)^21^ to generate alignment files of uniquely mapped reads with maximum allowed mismatch of 2 (-m 1 -n 2) for each alignment seed. ChIP-seq reads aligned to genome were extended to 150 bp from their 3’ end for further analysis. RNA-seq data were collected and single-end reads were aligned to human genome hg38 using TopHat (v2.1.0)^22^ with the parameters --min-segment-intron 50 --no-novel-indels --no-coverage-search, and only uniquely mapped reads were preserved.

### Identification of opened chromatin regions (OCR) by ATAC-seq

The ATAC-seq alignment files for CD127^high^ and CD127^low^ monocytes from three donors were used to call peaks for significantly opened chromatin regions using MACS2 (v2.1.1) (FDR < 0.05). The peaks from six samples were merged as total OCRs in LPS treated monocytes. To identify the differentially opened chromatin regions between CD127^high^ and CD127^low^ monocytes, the ATAC-seq fragments were counted in each OCR for each sample by FeatureCounts (v1.5.0)^23^. Subsequently, the fragments count was normalized to count per million mapped fragment for each sample. The normalized fragments count was used to identify differentially opened chromatin regions by edgeR (v3.28.1)^24^. Mean values of (fragments count+1) fold change (CD127^high^/CD127^low^) among three donors were log2 transformed, and differentially opened chromatin regions were identified by log2 transformed fold changes (CD127^high^/CD127^low^) ≥ 1 or ≤ −1 for CD127^high^ monocytes feature OCRs or CD127^low^ monocytes feature OCRs with cutoff of p-value < 0.05. To visualize ATAC-seq and ChIP-seq signals around open chromatin regions of interest, we firstly counted ATAC-seq fragments (-fragLength given) and ChIP-seq extended reads (-fragLength 150) every 10 bp from the center of OCR to ± 2.5 kb regions for each OCR by using annotatePeaks.pl program in HOMER (v4.7.2)^25^. The output counting matrices were used to generate average signals around OCRs by calculating the average fragments/reads count per bin (10 bp) per OCR.

### RNA-seq data analyses

For coverage of mapped RNA-seq reads in transcripts, the expression level of each gene transcript was calculated as normalized reads count per kilobase of transcript per million mapped reads (FPKM) using Cufflinks (v2.2.1)^26^. Differential gene expression between CD127^high^ and CD127^lwo^ monocytes from three donors was identified using DESeq2 (v1.27.9)^27^. Genes with p-value < 0.05 and mean (FPKM+1) fold changes (CD127^high^/CD127^low^) ≥ 1.5 or ≤ 0.67 among three donors were defined as highly expressed genes in CD127^high^ monocytes or highly expressed genes in CD127^low^ monocytes, respectively.

### scRNA-seq data analyses

For scRNA-seq of COVID-19 patients’ BALF cells, the Cell Ranger Software Suite (v.3.1.0) was used to perform sample de-multiplexing, barcode processing and single-cell 5′ unique molecular identifier (UMI) counting. Specifically, splicing-aware aligner STAR was used in FASTQs alignment. Cell barcodes were then determined based on the distribution of UMI counts automatically, and the gene-barcode matrices were saved for downstream analysis. In addition, one additional healthy control was acquired from the GEO database under accession number GSE128033. All samples were loaded as Seurat objects by using Seurat (v3.2.1)^14^, quality control for each cell were done with criteria as following: gene number between 200 and 6,000, UMI count > 1,000 and mitochondrial gene percentage < 0.1. All samples were further integrated to remove the batch effects with the parameter settings of the first 50 dimensions of canonical correlation analysis (CCA) and principal-component analysis (PCA). Integrated Seurat project was first normalized, and top 2,000 variable genes were then identified by using the Seurat analysis pipeline that has been described in our previous studies. Gene expression scaling and PCA was performed using the top 2,000 variable genes. Then UMAP was performed on the top 50 principal components for visualization, and graph-based clustering was simultaneously performed on the PCA-reduced data with the 1.2 resolution setting. According to the clustering result, 32 clusters were identified, and the annotations for each cluster were implemented based on the expression of marker genes that were used in our previous study^18^. Monocytes/macrophages (CD14^high^ CD68^high^) were extracted from the total BALF cells after annotation, and re-clustering (PCA and UMAP) was performed. The clusters showing expression of both monocyte/macrophage marker genes and T cell marker genes were excluded as doublets. Only the cells from COVID-19 patients were used for downstream analyses on both total BALF cells and monocytes/macrophages.

RA synovial scRNA-seq data sets^13^ were downloaded from ImmPort with the study accession code of SDY998. The reduction and clustering result from the original study were used. The monocytes clusters were extracted for downstream analysis.

The Cell Ranger Software Suite (v.3.1.0) was used to perform sample de-multiplexing, barcode processing and single-cell 5′ unique molecular identifier (UMI) counting, and gene-barcode matrices were generated for LPS-treated human CD14^+^ monocytes and RA patient’s PBMCs. The gene-barcode matrices were loaded as Seurat objects, and quality control for each cell was performed with criteria for LPS-treated human CD14^+^ monocytes (gene number between 200 and 4,500, UMI count > 1,000 and mitochondrial gene percentage < 0.15) and RA PBMCs (gene number between 200 and 6,000, UMI count > 1,000 and mitochondrial gene percentage < 0.1). After quality control, top 2,000 variable genes were identified, and gene expression scaling, PCA and UMAP clustering were performed for each data set. Marker genes for each cluster in each scRNA-seq data set were identified by using FindAllMarkers function in Seurat. According to the expression of well-studied PBMC marker genes, each cluster of RA PBMCs was annotated with certain cell type, and monocyte clusters were extracted for downstream analysis.

*IL7R*^+^ cells were identified based on the normalized expression (> 0) of *IL7R* for each cells in each scRNA-seq dataset. Inflammatory score was calculated based on the normalized average expression of eight inflammatory genes: *TNF, IL6, IL8, CCL2, CCL3, CCL4, CCL8* and *CXCL10*, which is implemented by AddModuleScore function in Seurat with 100 control for each inflammatory gene.

Integration of COVID-19 monocytes/macrophages, RA synovial monocytes and LPS-treated monocytes was implemented by a recently developed SCALEX method based on the original SCALE method^16^. Leiden clustering and UMAP visualization were performed based on the features extracted by SCALEX for integrated monocytes/macrophages, and marker genes for each cluster were identified. Correlation between clusters among three datasets were calculated as Pearson correlation coefficient, and the negative values were normalized to zero for heat map presentation.

### Statistical analysis

Types of statistical tests are indicated in figure legends. Statistical analyses were performed using GraphPad Prism Software (GraphPad Software Inc., La Jolla, CA, USA) for Student’s *t* test, and Wilcoxon rank-sum test was implemented using R (v.4.0.2). A value of *P* < 0.05 was considered statistically significant.

## Acknowledgements

We thank the patients and their families for their contribution to scientific research. We also thank clinical staffs and the COVID-19 Pathology Team from Third Military Medical University and Shanghai Jiao Tong University for performing the autopsy work. This research was supported by National Natural Science Foundation of China grants (31725010 and 31821003 to X.H), Emergency Project from Chongqing Health Commission (2020NCPZX01 to X.W.B.), Tsinghua University COVID-19 Scientific Research Program (2020Z99CFZ024 to X.H.), and funds from Tsinghua-Peking Center for Life Sciences (to X.H.).

## Author contributions

Bin Zhang performed experiments and bioinformatic analyses, analyzed and interpreted data, and wrote the manuscript. Yuan Zhang performed experiments and analyzed and interpreted data. Lei Xiong and Yuzhe Li assisted with integration of single cell RNA-seq datasets. Yunliang Zhang performed some of the monocyte experiments and assisted with single cell RNA-seq data analyses. Jiuliang Zhao, Hui Jiang, and Can Li assisted with collection of rheumatoid arthritis patient samples. Yunqi Liu performed STAT5 ChIP experiments. Xindong Liu, Haofei Liu, and Yi-Fang Ping performed pathological analyses of COVID-19 patient tissue samples. Qiangfeng Cliff Zhang directed the integration of single cell RNA-seq datasets and provided advices on bioinformatics. Zheng Zhang provided COVID-19 BALF single cell RNA-seq data sets and valuable advices. Xiu-Wu Bian directed pathological analyses of COVID-19 patient tissue samples. Yan Zhao provided rheumatoid arthritis patient samples and advices on the related experiments. Xiaoyu Hu conceptualized the project, supervised experiments, analyzed and interpreted data, and wrote the manuscript.

## Materials & Correspondence

Correspondence and requests for materials should be addressed to X.H.

## Data availability

Sequencing data sets are deposited in the Genome Expression Omnibus with assigned accession numbers as follows: RNA-seq and ATAC-seq in GSE159118, healthy donor scRNA-seq in GSE159113, RA scRNA-seq in GSE159117 and COVID-19 BALF scRNA-seq in GSE145926.

---

**Extended Data Figure 1.**
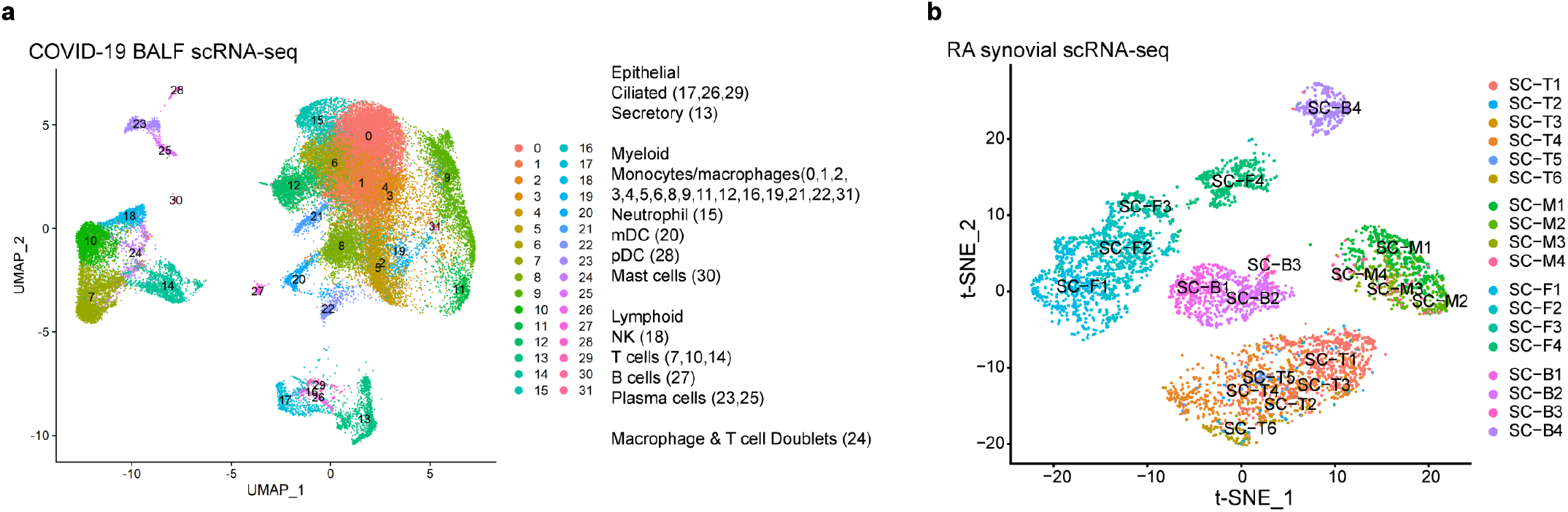
Clustering analyses of scRNA-seq data sets. UMAP projection of broncho-alveolar lavage fluid (BALF) cells from COVID-19 patients. Cell type annotations were labeled for each cluster. Monocyte/macrophage clusters (CD14^high^ CD68^high^) were used for the subsequent analyses. (**b**) t-SNE projection of synovial cells from RA patients. Cell type annotations were labeled for each cluster. T, M, F and B represent T cell, monocytes, fibroblasts and B cells, respectively. Monocyte clusters (CD14^high^) were used for the subsequent analyses.

**Extended Data Figure 2.**
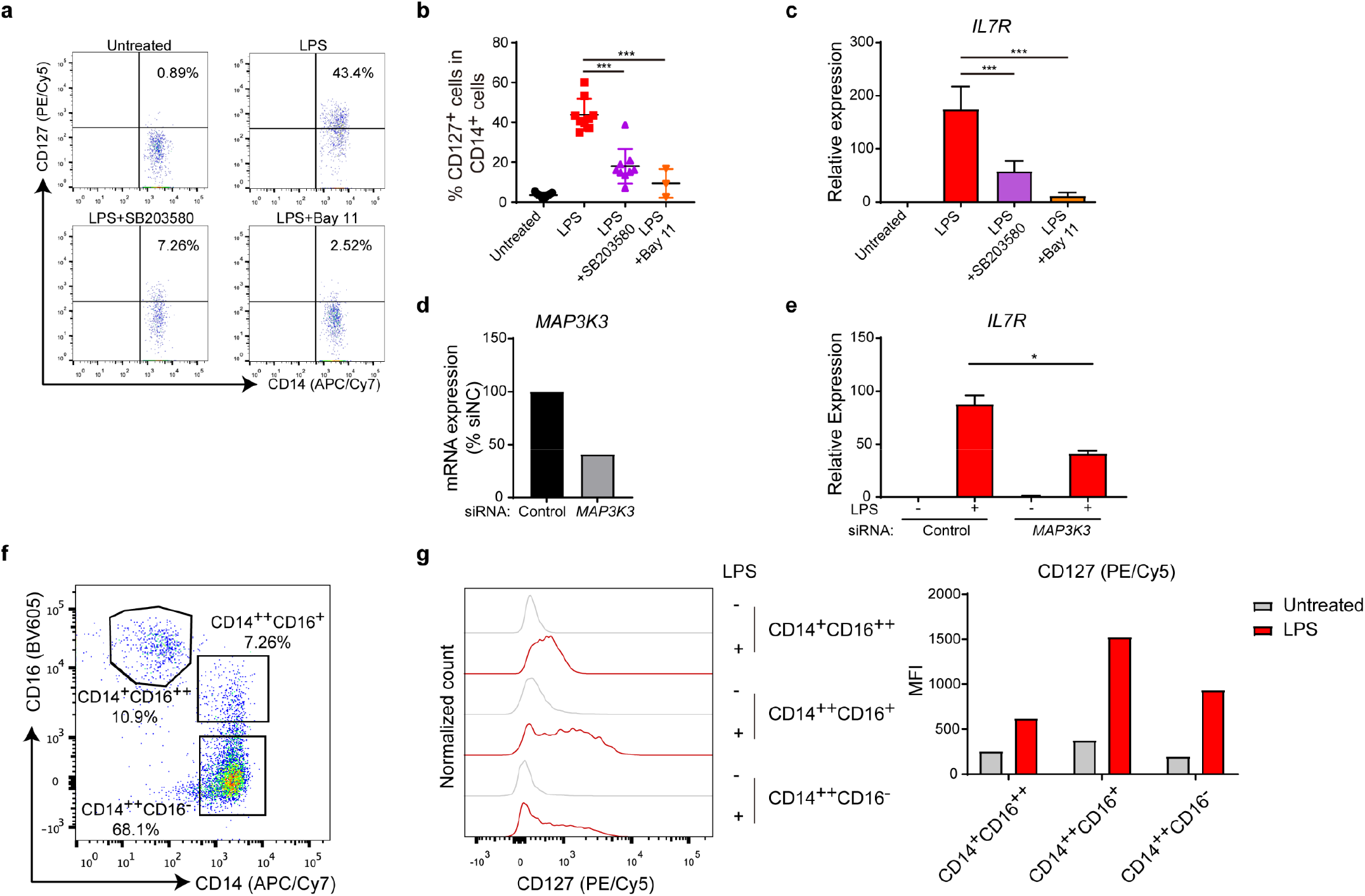
CD127 upregulation in activated human monocytes is dependent on canonical TLR signaling and observed in all monocyte subsets. (**a**, **b**) PBMCs form healthy donors were pretreated with DMSO or 10 μM SB203580 or 10 μM Bay 11-7082 (Bay 11) for 30 min and subsequently stimulated with or without 10 ng/ml LPS for 6 hours as indicated. The protein levels of CD127 were measured by flow cytometry and was shown as representative FACS distribution (**a**) and cumulative percentages (**b**) in CD14^+^ monocytes. Each data point represents the result from an independent experiment with cells obtained from a healthy donor. \*\*\**P*<0.001 by unpaired Student’s *t* test. Data are shown as means ± SD of multiple independent experiments as listed respectively: Untreated n = 9, LPS n = 9, SB203580 n = 9, Bay 11-7082 n = 3. (**c**) CD14^+^ monocytes were isolated form healthy donors’ PBMCs, and were pretreated with DMSO or 10 μM SB203580 or 10 μM Bay 11-7082 (Bay 11) for 30 min and subsequently stimulated with or without 10 ng/ml LPS for 6 h as indicated. The mRNA levels of *IL7R* were measured by qPCR. The relative expression was normalized to internal control (*GAPDH*) and expressed relative to the untreated sample. \*\*\**P*<0.001 by unpaired Student’s *t* test. Data are shown as means ± SD of multiple independent experiments as listed respectively: Untreated n = 9, LPS n = 9, SB203580 n = 6, Bay 11-7082 n = 3. (**d**, **e**) CD14^+^ monocytes were transfected with negative control or *MAP3K3* specific short interfering RNAs. Two days post transfection, cells were stimulated with LPS (10 ng/ml) for 3 h. Knock down efficiency of *MAP3K3* was examined (**d**), and mRNA induction of *IL7R* by LPS stimulation in siControl and si*MAP3K3* transfected cells was measured by qPCR (**e**). Relative expression was normalized to internal control (*GAPDH*) and expressed relative to LPS untreated siControl sample. \**P*<0.05 by paired Student’s *t* test. *IL7R* expression data are shown as means ± SD of three independent experiments with cells from one healthy donor for each data set. (**f**, **g**) Three monocyte populations were gated by CD14 and CD16 expression in flow cytometry analysis of human PBMCs. CD127 was also stained in PBMCs with or without LPS stimulation, and CD127 upregulation in three human monocyte populations was analyzed by gating strategy shown in (**f**). Representative FACS distribution (**g**, left) and mean fluorescence intensity (MFI) (**g**, right) for CD127 expression are shown.

**Extended Data Figure 3.**
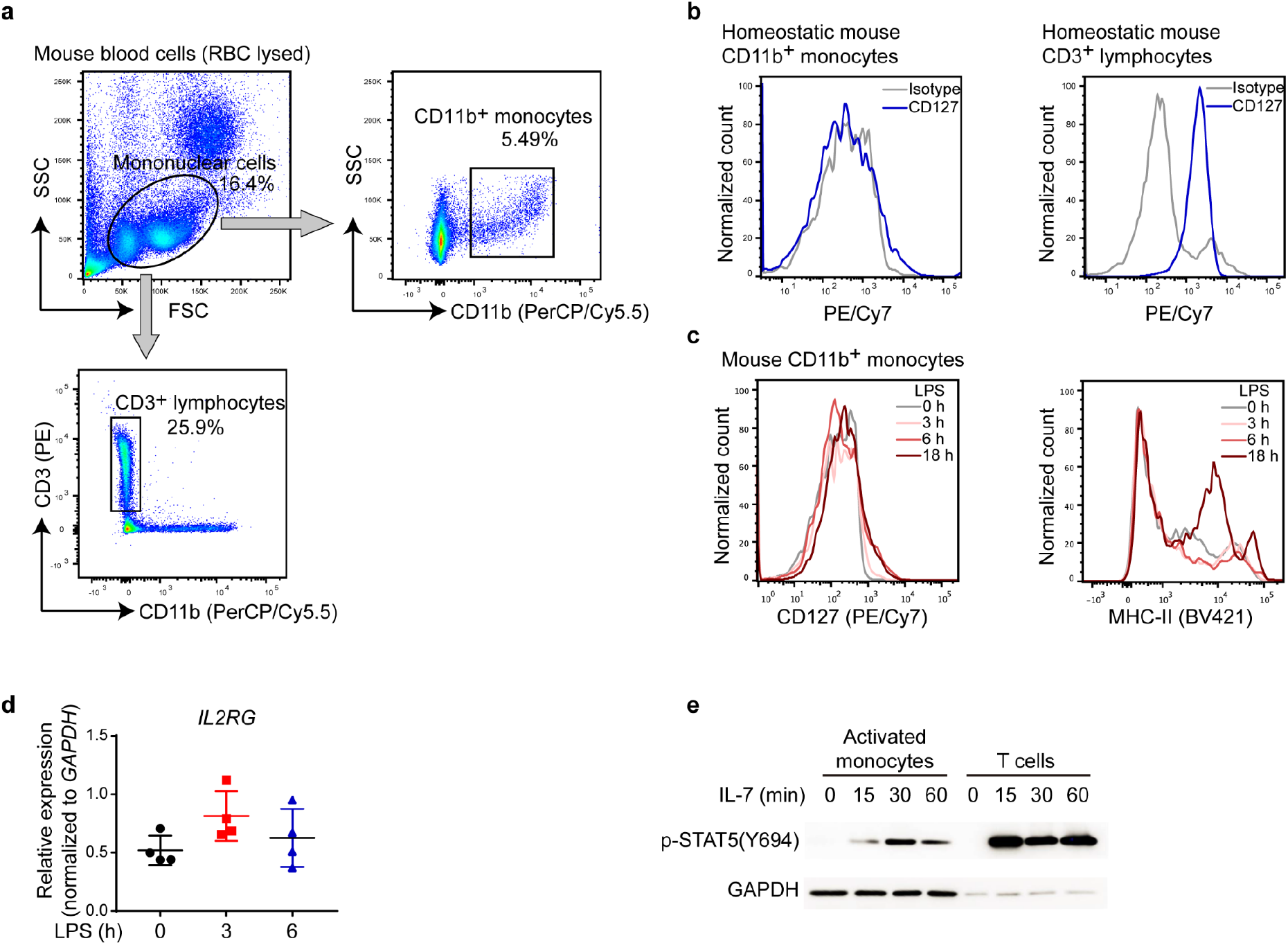
Human but not mouse activated monocytes are CD127 positive and functionally competent for IL-7 receptor signaling. (**a**) Gating strategy for CD11b^+^ mouse monocytes and CD3^+^ mouse lymphocytes from red blood cell (RBC) lysed mouse peripheral blood cells. (**b**, **c**) Mouse blood cells (RBC lysed) were treated with or without 100 ng/ml LPS for different time points as indicated. The expression of CD127 (**b**) and MHC-II (**c**) on mouse CD11b^+^ monocytes was analyzed by flow cytometry. (**d**) Human CD14^+^ monocytes were treated with 10 ng/ml LPS for 3 h and 6 h, the mRNA levels of *IL2RG* were measured by qPCR. Each data point represents the result from an independent experiment with cells obtained from a healthy donor. Relative expression was normalized to *GAPDH* as internal control. Data are shown as means ± SD of four independent experiments. (**e**) Human CD14^+^ monocytes were pretreated with 10 ng/ml LPS for 6 h, and subsequently stimulated with recombinant human IL-7 (10 ng/ml) for the indicated time. Meanwhile, CD3^+^ T cells from the same donor were treated with 10 ng/ml IL-7 for the indicated time. The protein levels of p-STAT5(Y694) were detected by western blotting. GAPDH was used as a loading control.

**Extended Data Figure 4.**
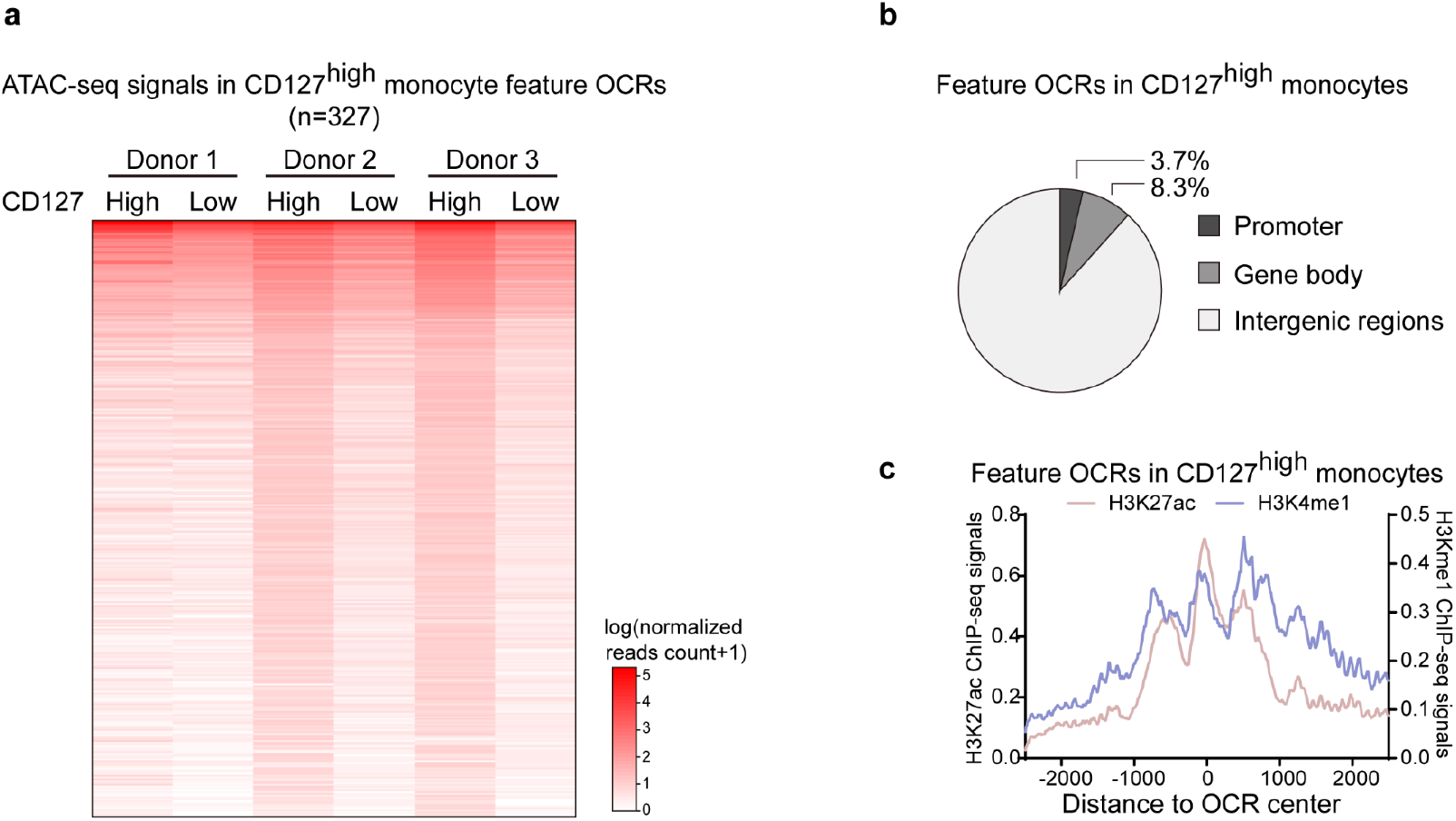
Open chromatin regions (OCRs) in CD127high monocytes display enhancer-like features. (**a**) Heat map shows the ATAC-seq signals in OCRs featured in CD127^high^ monocytes and in CD127^low^ monocytes from three independent experiments with three blood donors. The ATAC-seq reads count was normalized (per 10 million mapped reads) and then log10(n + 1) transformed for expression in heat map. (**b**) Pie graph shows the genomic distribution of CD127^high^ monocyte-featured OCRs. (**c**) Average ChIP-seq signals of H3K27ac and H3K4me1 in LPS activated monocytes were assessed around CD127^high^ monocyte-featured OCRs and expressed by different colors as indicated in the plot.

**Extended Data Figure 5.**
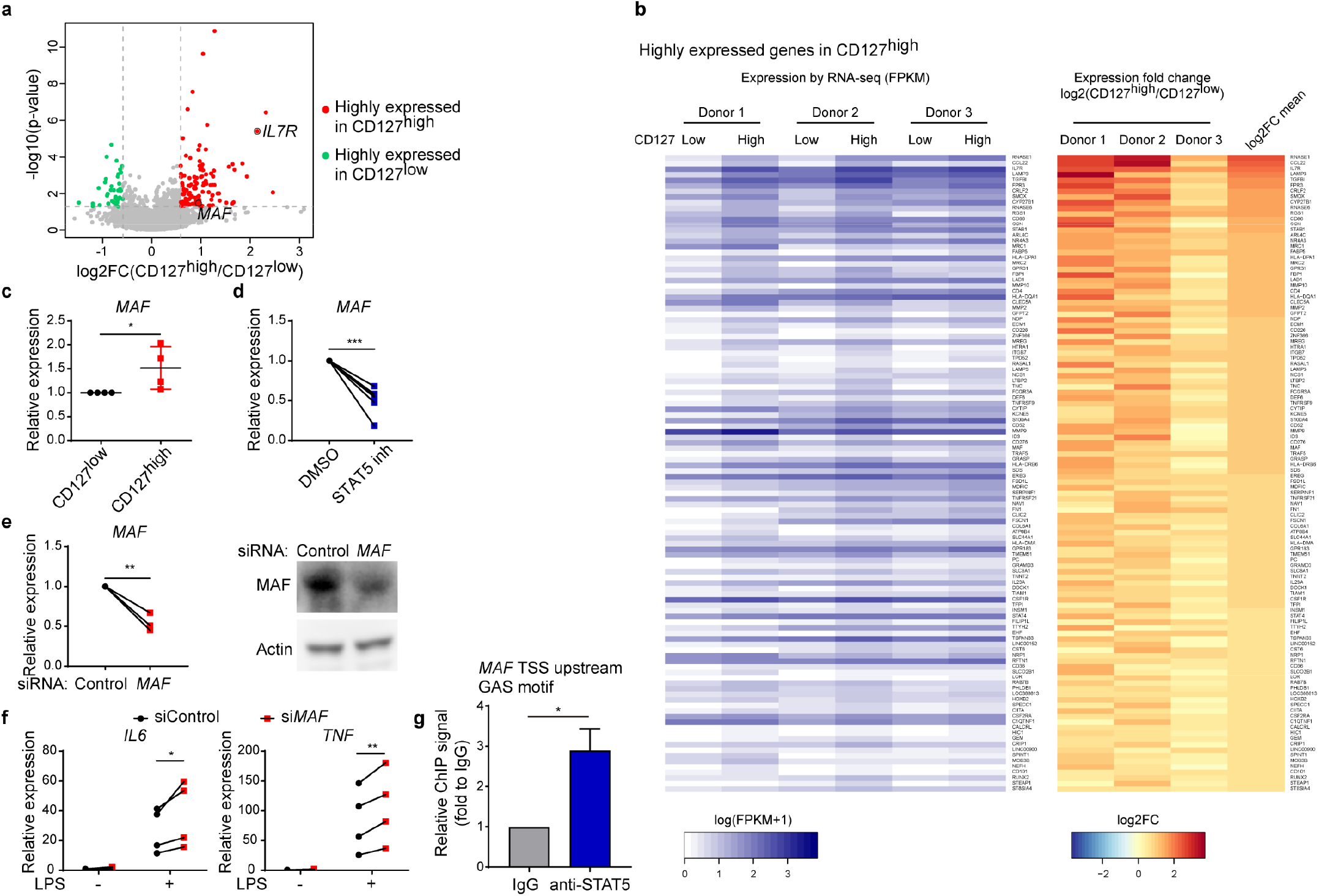
CD127-STAT5-c-Maf axis imposes an anti-inflammatory loop in activated monocytes. (**a**) Volcano plot shows the differentially expressed genes by RNA-seq between CD127^high^ and CD127^low^ populations from CD14^+^ monocytes upon 6 h LPS stimulation. Red colors indicate highly expressed genes in CD127^high^ population, and green colors indicate highly expressed genes in CD127^low^ population. Two CD127^high^ highly expressed genes, *IL7R* and *MAF*, were highlighted with black circles. The dash lines indicate the threshold of p-value (p-value < 0.05) and fold change (≥1.5 fold for upregulation and ≤0.67 fold for downregulation) for identifying differentially expressed genes. Fold changes are shown as mean value of CD127^high^/ CD127^low^ ratio from three healthy donors. (**b**) Heat map shows the highly expressed genes in CD127^high^ monocytes in (A). The original expression (log10 transformed FPKM+1, left) and expression fold change (log2 transformed CD127^high^/CD127^low^, right) were shown. Fold changes are shown as the individual value in each donor and the mean value from three donors. (**c**) The mRNA expression of *MAF* in CD127^high^ and CD127^low^ monocytes was measured by qPCR. Relative expression was normalized to internal control (*GAPDH*) and expressed relative to CD127^low^ sample. Each data point represents the result from an independent experiment with cells obtained from a healthy donor. \**P*<0.05 by paired Student’s *t* test. Data are shown as means ± SD of four independent experiments. (**d**) CD14^+^ monocytes were pretreated with STAT5 inhibitor (100 μM) for 2 h and then were stimulated with LPS (10 ng/ml) for 6 h. mRNA of *MAF* was measured by qPCR. Relative expression is normalized to internal control (*GAPDH*) and expressed relative to DMSO treated sample. Each data point represents the result from an independent experiment with cells obtained from a healthy donor. \*\*\**P*<0.001 by paired Student’s *t* test. The results from six independent experiments are shown. (**e**, **f**) CD14^+^ monocytes were transfected with negative control or *MAF* specific short interfering RNA (siControl or si*MAF*). Two days post transfection, knockdown efficiency was assessed by measuring *MAF* mRNA with qPCR (**e**left, relative expression is normalized to internal control (*GAPDH*) and expressed relative to siControl group) and MAF protein levels with immunoblotting (**e**right). Two days post transfection, cells were stimulated with LPS (10 ng/ml) for 3 h and mRNA levels of *IL6* and *TNF* were measured by qPCR (**f**). Relative expression was normalized to internal control (*GAPDH*) and expressed relative to LPS untreated siControl sample. Each data point represents the result from an independent experiment with cells obtained from a healthy donor. \**P*<0.05, \*\**P*<0.01 by paired Student’s *t* test. The results from four independent experiments are shown. (**g**) Occupancy of STAT5 on the *MAF* transcription start site (TSS) upstream GAS motif was assessed by ChIP-qPCR in THP-1 cells with 6 h LPS stimulation prior to 30 min IL-7 stimulation. IgG ChIP was used as negative control, and relative STAT5 ChIP signals were expressed relative to IgG ChIP control sample. \**P*<0.05 by paired Student’s *t* test. Data are shown as means ± SD of three independent experiments.

**Extended Data Figure 6.**
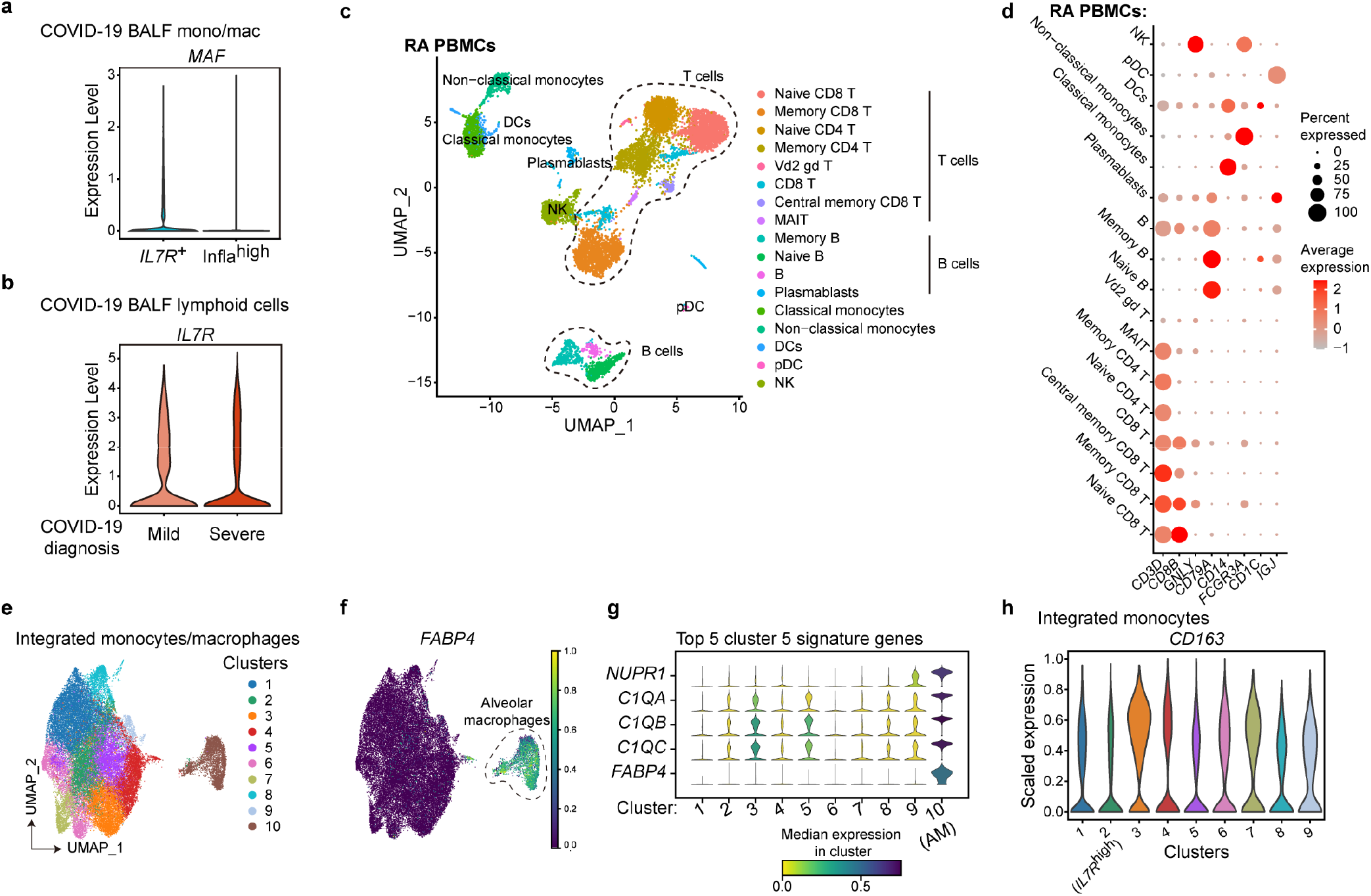
CD127 expression designates a functionally distinct monocyte subset. (**a**) Violin plot shows the *MAF* expression distribution in *IL7R*^+^ cells and infla^high^ cells in monocytes/macrophages from COVID-19 patients’ BALF in Fig. 4a. (**b**) Lymphoid cells from COVID-19 patients’ BALF were subgrouped by the diagnosed disease severity. Violin plot shows the expression of *IL7R* in BALF lymphoid cells from mild or severe COVID-19 patients. (**c**) UMAP projection of PBMCs from RA patient. Cell type annotations were labeled for each cluster. (**d**) Heat map shows the expression of hallmark genes in different cell clusters from RA PBMCs. The scaled average expression levels of marker genes and percentage of cell expressing marker genes were expressed by color and size of each dot corresponding to cell clusters, respectively. (**e**) UMAP projection of integrated monocytes/macrophages from COVID-19 BALF, RA synovial cavity and LPS-activated monocytes. Ten clusters by unsupervised clustering were indicated by different colors in plot. (**f**) UMAP projection of integrated monocytes/macrophages in (**e**). *FABP4* expression in cells was quantitatively visualized by the indicated colors. Cells corresponding to Cluster 5 were highlighted by dotted line as alveolar macrophages given the specific *FABP4* expression pattern. (**g**) The stacked violin plot shows the expression of top 5 cluster 5 signature genes in all ten clusters shown in (**e**). Median expression levels for each gene in each cluster were expressed by colors as indicated. (**h**) Violin plot shows the expression of *CD163* among nine clusters of integrated monocytes in Fig. 4f, in which cluster 2 was designated as *IL7R*^high^ cluster.

**Extended Data Figure 7.**
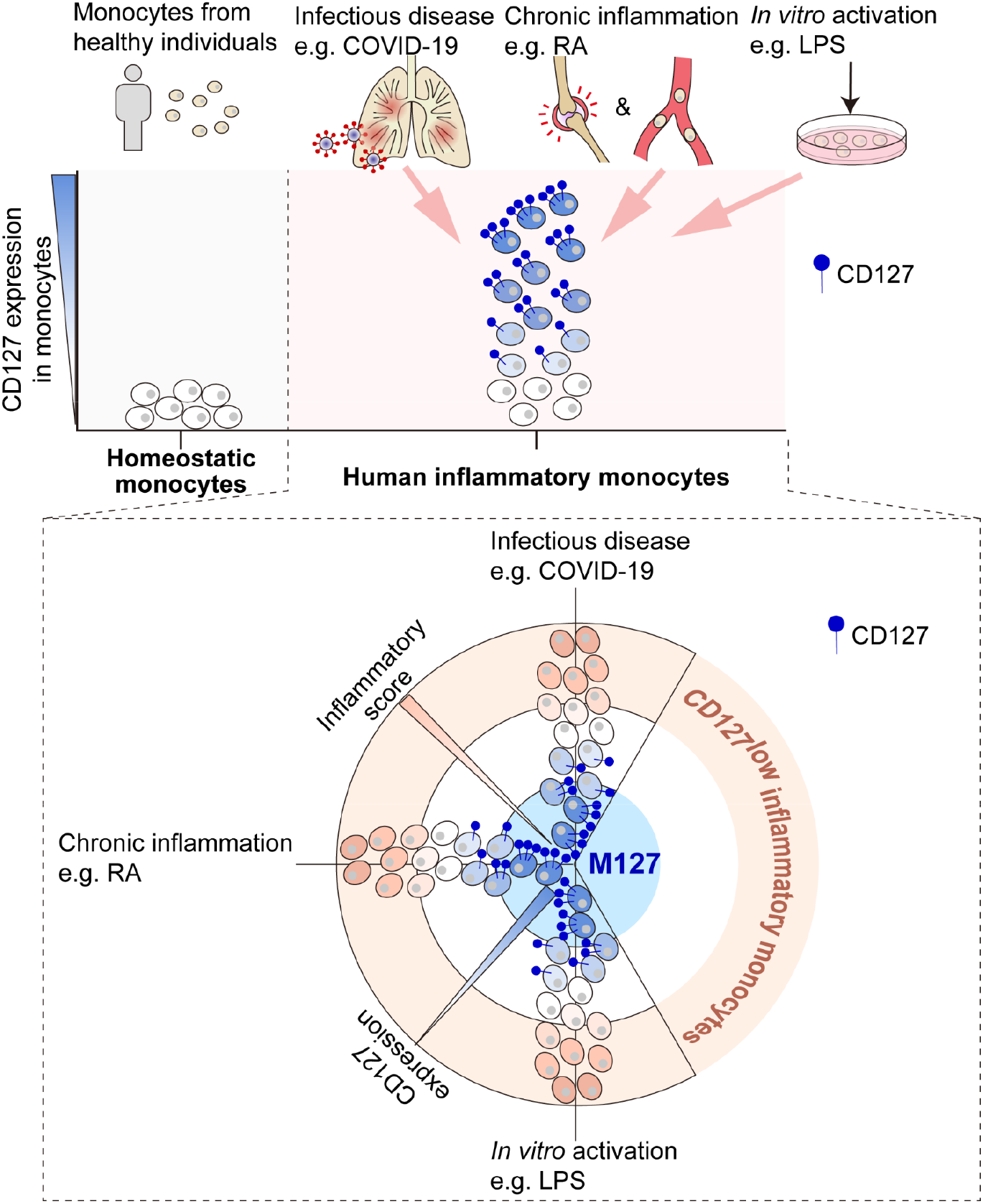
M127 represents a unique hypo-inflammatory monocyte population in human inflammatory diseases. CD127 is minimally expressed on homeostatic monocytes from healthy individuals. Upregulation of CD127 is a unified hallmark event for human monocyte activation under various inflammatory conditions including COVID-19, RA and stimulated blood monocytes. CD127^+^ inflammatory monocytes, M127, display unique gene signatures and subdued inflammatory phenotypes imposed by CD127-STAT5 signaling. As a result, CD127 levels inversely correlate with the inflammatory capacity of human monocytes. Identification and characterization of M127 highlight the previously underappreciated functional diversity among human monocytes in inflammatory diseases.

**Extended Data Table 1.**
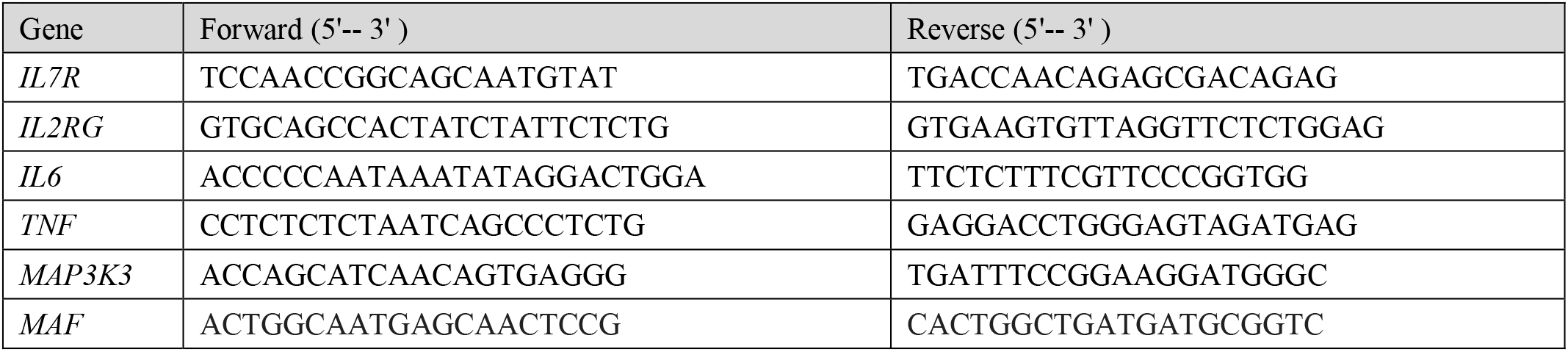
qPCR primers.

**Extended Data Table 2.** Antibodies used for the experiments.

| Antibodies | Manufacturer | Catalog number |
| --- | --- | --- |
| Anti-CD68 antibody | Abcam | ab201340 |
| Anti-human CD127 antibody | Invitrogen | PA5-97870 |
| Goat anti-rabbit IgG (H&L)-HRP conjugated | Bioeasy (Beijing) Technology CO. Ltd | BE0101 |
| Goat anti-mouse IgG (H&L)-HRP conjugated | Bioeasy (Beijing) Technology CO. Ltd | BE0102 |
| Alexa Fluor™ 488 goat anti-mouse IgG | Life Technologies | B40941 |
| Alexa Fluor™ 555 goat anti-rabbit IgG | Invitrogen | A27039 |
| Anti-mouse CD3ε antibody PE | Biolegend | 100307 |
| Anti-mouse CD11b PerCP-Cyanine5.5 | eBioscience | 45-0112-82 |
| Anti-mouse CD127 PE-Cyanine7 | Biolegend | 135013 |
| PE-Cy7 Rat IgG2a, $\kappa$ isotype control antibody | Biolegend | 400521 |
| Anti-mouse MHC class II BV421 | eBioscience | 48-5321-82 |
| Anti-human CD127 antibody PE-Cyanine5 | Biolegend | 351323 |
| Anti-human CD14 antibody APC-Cyanine7 | Biolegend | 301820 |
| Anti-human IL-6 antibody PE | Biolegend | 501106 |
| Anti-human TNF-α antibody PE | Biolegend | 502908 |
| PE Rat IgG2b, $\kappa$ isotype control antibody | Biolegend | 400607 |
| Anti-human CD16 antibody BV605 | Biolegend | 302039 |
| Anti-GAPDH Mouse Monoclonal Antibody | Bioeasy (Beijing) Technology CO. Ltd | BE0023 |
| Anti-β-Actin Mouse Monoclonal Antibody | Abclonal | AC026 |
| Anti-Phospho-Stat5 (Tyr694) Rabbit Monoclonal Antibody | Cell Signaling Technology | 9314S |
| Anti-Stat5 Antibody | Cell Signaling Technology | 9363S |
| Anti-c-Maf Antibody | Santa Cruz Biotechnology | sc-518062 |

## References

1 Merad, M. & Martin, J. C. Pathological inflammation in patients with COVID-19: a key role for monocytes and macrophages. Nature reviews. Immunology 20, 355–362 (2020).

2 Donlin, L. T. et al. Insights into rheumatic diseases from next-generation sequencing. Nat Rev Rheumatol 15, 327–339 (2019).

3 Ermann, J., Rao, D. A., Teslovich, N. C., Brenner, M. B. & Raychaudhuri, S. Immune cell profiling to guide therapeutic decisions in rheumatic diseases. Nat Rev Rheumatol 11, 541–551 (2015).

4 Shi, C. & Pamer, E. G. Monocyte recruitment during infection and inflammation. Nature reviews. Immunology 11, 762–774 (2011).

5 van der Poll, T., van de Veerdonk, F. L., Scicluna, B. P. & Netea, M. G. The immunopathology of sepsis and potential therapeutic targets. Nature reviews. Immunology 17, 407–420 (2017).

6 Mildner, A., Yona, S. & Jung, S. A close encounter of the third kind: monocyte-derived cells. Adv Immunol 120, 69–103 (2013).

7 Auffray, C., Sieweke, M. H. & Geissmann, F. Blood monocytes: development, heterogeneity, and relationship with dendritic cells. Annu Rev Immunol 27, 669–692 (2009).

8 Guilliams, M., Mildner, A. & Yona, S. Developmental and Functional Heterogeneity of Monocytes. Immunity 49, 595–613 (2018).

9 Nathan, C. & Ding, A. Nonresolving inflammation. Cell 140, 871–882 (2010).

10 Vabret, N. et al. Immunology of COVID-19: Current State of the Science. Immunity 52, 910–941 (2020).

11 Barata, J. T., Durum, S. K. & Seddon, B. Flip the coin: IL-7 and IL-7R in health and disease. Nat Immunol 20, 1584–1593 (2019).

12 Kalliolias, G. D. & Ivashkiv, L. B. TNF biology, pathogenic mechanisms and emerging therapeutic strategies. Nat Rev Rheumatol 12, 49–62 (2016).

13 Zhang, F. et al. Defining inflammatory cell states in rheumatoid arthritis joint synovial tissues by integrating single-cell transcriptomics and mass cytometry. Nat Immunol 20, 928–942 (2019).

14 Stuart, T. et al. Comprehensive Integration of Single-Cell Data. Cell 177, 1888–1902 e1821 (2019).

15 Smale, S. T. & Natoli, G. Transcriptional control of inflammatory responses. Cold Spring Harb Perspect Biol 6, a016261 (2014).

16 Xiong, L. et al. SCALE method for single-cell ATAC-seq analysis via latent feature extraction. Nat Commun 10, 4576 (2019).

17 Ronit, A. et al. Compartmental immunophenotyping in COVID-19 ARDS: a case series. The Journal of allergy and clinical immunology 10.1016/j.jaci.2020.09.009 (2020).

18 Liao, M. et al. Single-cell landscape of bronchoalveolar immune cells in patients with COVID-19. Nat Med 26, 842–844 (2020).

19 Novakovic, B. et al. beta-Glucan Reverses the Epigenetic State of LPS-Induced Immunological Tolerance. Cell 167, 1354–1368 e1314 (2016).

20 Langmead, B. & Salzberg, S. L. Fast gapped-read alignment with Bowtie 2. Nat Methods 9, 357–359 (2012).

21 Langmead, B., Trapnell, C., Pop, M. & Salzberg, S. L. Ultrafast and memory-efficient alignment of short DNA sequences to the human genome. Genome Biol 10, R25 (2009).

22 Kim, D. et al. TopHat2: accurate alignment of transcriptomes in the presence of insertions, deletions and gene fusions. Genome Biol 14, R36 (2013).

23 Liao, Y., Smyth, G. K. & Shi, W. featureCounts: an efficient general purpose program for assigning sequence reads to genomic features. Bioinformatics 30, 923–930 (2014).

24 Robinson, M. D., McCarthy, D. J. & Smyth, G. K. edgeR: a Bioconductor package for differential expression analysis of digital gene expression data. Bioinformatics 26, 139–140 (2010).

25 Heinz, S. et al. Simple combinations of lineage-determining transcription factors prime cis-regulatory elements required for macrophage and B cell identities. Mol Cell 38, 576–589 (2010).

26 Trapnell, C. et al. Differential gene and transcript expression analysis of RNA-seq experiments with TopHat and Cufflinks. Nat Protoc 7, 562–578 (2012).

27 Love, M. I., Huber, W. & Anders, S. Moderated estimation of fold change and dispersion for RNA-seq data with DESeq2. Genome Biol 15, 550 (2014).

